# Molecularly defined auditory neuron subtypes show different vulnerabilities to noise- and age-related synaptopathy in mice

**DOI:** 10.1101/2025.08.27.672747

**Authors:** Joy A. Franco, Taylor G. Copeland, Ryan D. Merrow, Lisa V. Goodrich

## Abstract

Neuronal subtype-specific synaptopathy is a hallmark of many forms of neurodegeneration. We examined the cellular basis for synaptic vulnerability in the auditory system, where three subtypes of spiral ganglion neurons (SGNs)—Ia, Ib, and Ic—carry acoustic information from the cochlea to the brain. In response to noise and aging, a subset of synapses between inner hair cells and SGNs are lost, but it is unclear how this loss varies across SGN subtypes. Using genetic labelling, we showed that Ia SGNs have larger post-synaptic densities (PSDs) than Ib and Ic SGNs and are the most resilient subtype. Ia PSD volumes increased with age and were unchanged after noise exposure. By contrast, average Ib/Ic PSD volumes did not change with age but decreased with noise. Genetic reprogramming of Ib/Ic neurons to a Ia-like identity provided significant protection against noise-induced synaptopathy, linking identity to resilience and providing an entry point for therapeutics.

---

Excitatory, glutamatergic synapses mediate the rapid transmission and faithful encoding of information between cells. However, when too much glutamate is released, synapses may suffer excitotoxic damage and be eliminated, resulting in synaptopathy^1^. Synaptic dysfunction and loss precede neuronal cell death in a range of neurodegenerative disorders^2^. Despite the ubiquitous presence of excitatory synapses, some synapses are vulnerable and others are resilient, exemplified by the selective vulnerability of certain regions of the brain in Alzheimer’s Disease^3^. What makes some synapses better able to withstand excitotoxic insults remains unknown.

Type I spiral ganglion neurons (SGNs) of the auditory system provide a tractable opportunity for uncovering the relationship between a neuron’s genetic identity and the vulnerability of its synapses. Sound detection begins in the cochlea, where inner hair cells (IHCs) respond to sound via mechanically sensitive ion channels. IHCs then release glutamate from presynaptic ribbon structures onto the postsynaptic terminals of Type I SGNs, which fire action potentials that travel from the cochlea to the brain. Excessively intense stimuli lead to loss of some IHC-SGN synapses, likely due to glutamate excitotoxicity^4,5^. Although uniformly glutamatergic, individual IHC-SGN synapses exhibit a range of morphologies and properties that correlate with anatomical and physiological differences among the SGN subtypes^6^. Type I SGN subtypes were initially defined by differences in their spontaneous firing rates (SR) and the minimum sound intensity required for action potential generation^7^, with low SR fibers innervating the modiolus-facing (modiolar) region of the IHC and high SR fibers innervating the opposing, pillar side. Type I SGNs were subsequently divided into molecularly distinct Ia, Ib, and Ic SGNs that differ in expression of many synaptic and calcium signaling proteins^8–10^. Based on the position of their terminals on the IHC and their firing patterns *in vitro*, Ia SGNs correspond best to high SR SGNs, whereas Ic SGNs are likely low SR SGNs^11^ (Fig. 1a). In contrast to the complexity of synaptic connectivity in the central nervous system (CNS), Type I SGNs receive inputs from only one or two IHCs and each IHC signals to 20 or fewer Type I SGNs^12^, making it straightforward to quantify and assess synaptic properties in a comprehensive manner. Additionally, the efficacy of signaling through IHC-SGN synapses can be assessed by recording auditory brainstem responses (ABRs), since the amplitude of the first coordinated neuronal response correlates with the number of cochlear synapses^13^. Understanding why some IHC-SGN synapses are lost while others persist offers a valuable chance to characterize mechanisms of selective vulnerability among neurons that are otherwise very similar to each other.

**Fig. 1:**
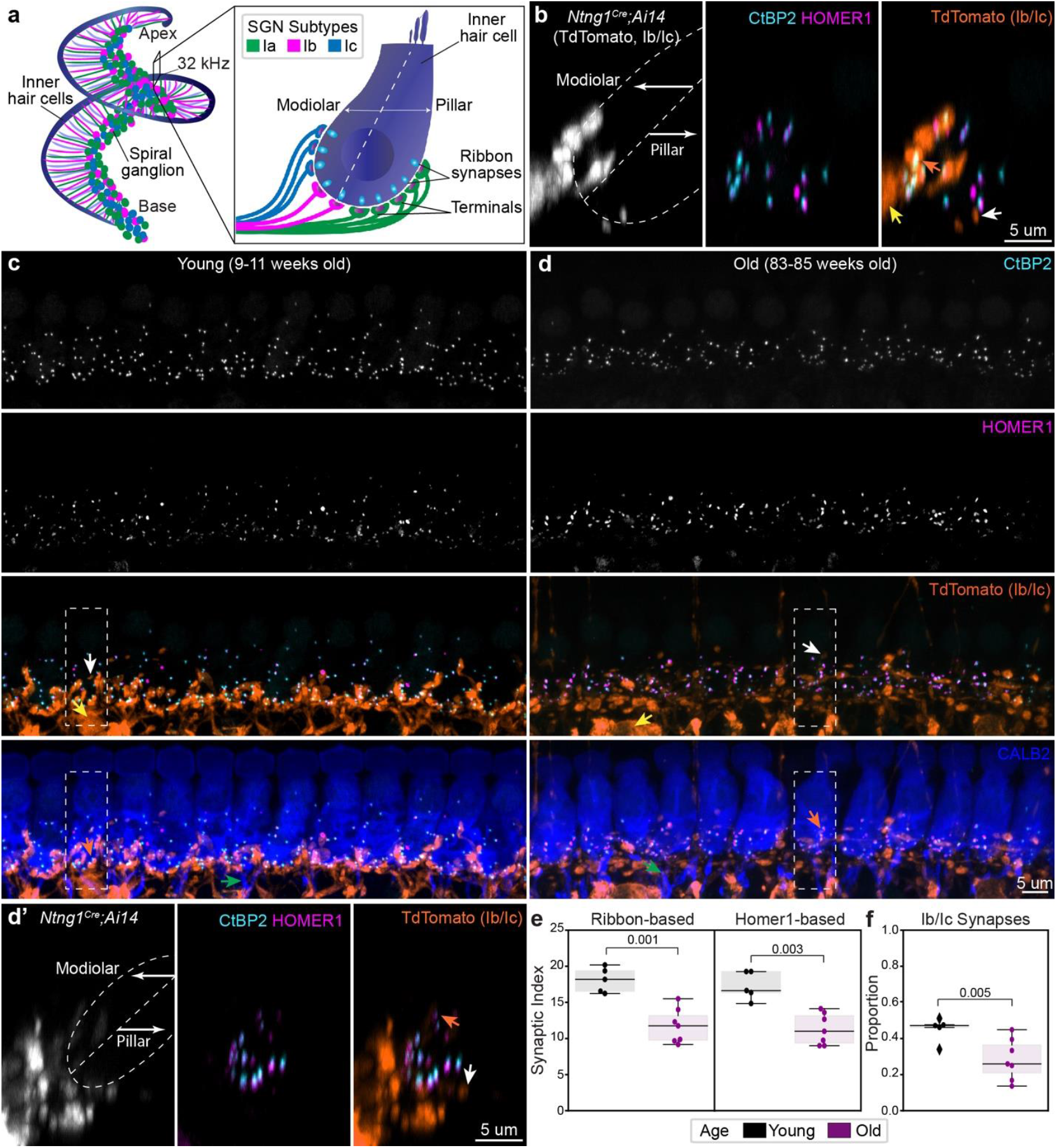
Type Ib/Ic SGNs are selectively vulnerable to age-related synaptopathy. **a** Schematic representation of SGN innervation of IHCs in the cochlea, color-coded by molecularly defined neuronal subtype. **b** Orthogonal view of an IHC in *Ntng1*^*Cre*^*;Ai14* mouse, where Type Ib/Ic SGN terminals (tdTomato+) primarily innervate the modiolar face of the IHC, overlapping with a subset of post-synaptic markers. **c** and **d** Representative high-magnification micrographs used for synapse and terminal analysis from young (**c**, 9-11 weeks) and old animals (**d**, 83-85 weeks). Dashed outlines correspond to regions used to generate maximum-intensity projections of virtually sliced z-stacks shown in **b** and **d’. e** Quantification of synaptic density using either ribbon-based or Homer1-based approaches. **f** Proportion of Ib/Ic synapses as quantified from Homer1-based analysis. For **e** and **f**, each dot corresponds to a single field of view (one per animal), boxes represent interquartile ranges, horizontal lines represent the median, box height represents the interquartile range (IQR), and whiskers extend to data points farthest from the median but within 1.5x the IQR from the box edge. Diamonds mark data points outside whiskers, classified as potential outliers by Python Seaborn package. Statistical analyses performed with one-sided Mann-Whitney tests for all and include data points classified as potential outliers. Arrows for all panels: orange=Type Ib/Ic synapses, green=fibers with highest levels of CALB2, yellow=glial-like cells, white = efferents. Multichannel overlay for **b** and **d’** with CALB2 to show hair cell morphology is provided in Supplementary Fig. 1e.

Although cochlear synaptopathy does not affect all SGNs equally, it remains unclear which SGN subtypes are most resilient and which are most vulnerable. Noise exposure and aging can both lead to cochlear synaptopathy that has physiological relevance to hearing loss in humans. In humans and mice, early life exposure to excessive noise is a strong predictor of age-related hearing loss and is correlated with a permanent loss of synapses and eventual SGN death. In humans, this pathology is thought to underlie “hidden hearing loss” where auditory thresholds are maintained but patients report hearing difficulties in noisy environments^14–17^. SGN synaptic heterogeneity is thought to endow the cochlea with a broad dynamic range such that high SR fibers encode low intensity auditory information and low SR fibers are recruited at louder sound intensities, when high SR fibers are saturated. Loss of low SR SGNs, would in theory reduce the dynamic range of the cochlea, making it more difficult to amplify quiet sounds in noisy landscapes. However, it remains unclear whether low SR SGNs are indeed those most impacted by synaptopathy and as a result, the full importance of SGN synaptic heterogeneity to auditory function has yet to be elucidated. Being able to genetically identify the most vulnerable IHC-SGN synapses would provide a handle for directly probing this relationship.

Physiological and anatomical approaches used in prior studies of synaptic resilience have faced several technical limitations. *In vivo* recordings of the auditory nerve in noise exposed guinea pigs showed a reduction in the presence of low SR SGNs^18^, which likely correspond to Ic SGNs^11^. However, no firm conclusions could be made about the identity of the affected neurons, as it was unclear whether the change in firing properties was due to a loss of specific synapses or because of noise-induced alterations in the properties of the remaining synapses and/or the post-synaptic neuron. Additionally, similar experiments did not observe a loss of low SR fibers in noise exposed mice and instead showed increased firing rates in some cases^19^. Anatomical studies of the noise exposed cochlea also suggested Ic SGNs are more vulnerable, as synapses on the modiolar side (where Ib and Ic fibers are positioned) were preferentially lost. Low SR SGNs, presumably Ic neurons, also seem to be lost from the aged cochlea, suggested both by *in vivo* recordings in aged gerbils^20^ and the fact that the surviving synapses in older animals primarily reside on the pillar side of the IHC^21^, suggesting they belong to Ia SGNs. *In situ* hybridization for Ia and Ic SGN markers similarly showed that Ia SGNs survive into old age whereas Ic SGNs are lost^8^. Since SGN loss can be exacerbated by early life exposure to noise^14^, the most parsimonious interpretation is that aging Ic SGNs lose their synapses and then die. However, the pattern of synapse loss along the pillar-modiolar axis is not consistent across the multiple strains of mice examined^22^, so it is also possible that different mechanisms are involved. Additionally, synaptic position is not an entirely reliable indicator of Type I SGN subtypes, as positions can shift during repair^23^. Thus, the question of whether molecularly defined SGN subtypes show different vulnerabilities to noise-induced or age-related synaptopathy remains unanswered.

Here, we directly evaluated subtype vulnerability by using genetic approaches to label Ia, Ib, and Ic SGNs. Our data confirm that Ic subtypes are the most vulnerable to both age-related and noise-induced synaptopathy. Furthermore, a genetic manipulation that shifts Ic SGNs towards Ia identities without changing the position of their synapses resulted in enhanced resilience to noise exposure. Together, these findings link genetic identity to resilience, offering a fruitful platform for identifying relevant molecular programs and developing targeted therapeutics in the future.

## Results

### Type Ia SGNs are most resilient to age-related synaptopathy

To track synapse loss from molecularly defined SGN subtypes, Ib and Ic SGNs were genetically labeled using *Netring1*^Cre/+^;*Rosa26*^Ai14+^ mice (*Ntng1*^*Cre*^*;Ai14*). Consistent with the enrichment of *Netring1* (*Ntng1*) in adult Ib and Ic SGNs^8^, tdTomato is expressed in the vast majority of Ib and Ic SGNs and not in Ia SGNs in *Ntng1*^*Cre*^*;Ai14* mice^24^. Cytosolic tdTomato fills the Ib and Ic SGN peripheral processes and their terminals, which were mostly on the modiolar side of the IHC (Fig. 1b), though this boundary was not strictly observed. Since this approach does not allow us to distinguish Ib’s from Ic’s, we refer to these co-labelled SGNs collectively as “Ib/Ic.” IHC-SGN synapses were identified with antibodies for CtBP2, which detects Ribeye in the pre-synaptic ribbons, and Homer1, which marks the post-synaptic densities (PSDs) in the SGN terminals (Fig. 1b-d). Each PSD could be reliably scored as belonging to Ib/Ic (tdTomato+, Fig. 1b-d orange arrows) or Ia (tdTomato-) SGN subtypes by assessing the degree of overlap between Homer1 and tdTomato. TdTomato-peripheral processes expressed Calretinin (CALB2), confirming that they came from Ia SGNs (Fig. 1c-d, green arrows). TdTomato was also visible in efferents (Fig. 1b-d, white arrows) and glial cells (Fig. 1, yellow arrows). However, these structures are spatially and morphologically distinct from Homer1 containing terminals and therefore did not confound analysis.

Visualization of synapses in cochleae from young (9-11 weeks) and old (83-85 weeks) *Ntng1*^*Cre*^*;Ai14* animals confirmed age-related synaptopathy. For each animal, a high-magnification, 100 um wide field of view was acquired at the 32 kHz region, which included on average 11.4+/-0.4 IHCs in young animals (N=5) and 7.4+/-0.6 IHCs in old animals (N=7). CtBP2+ and Homer1+ puncta density within the boundaries of an IHC was reduced in old animals when compared to young controls (Fig. 1c, d). In addition, fewer Ib/Ic terminals were apparent on and under IHCs in the old animals (Fig. 1c’, d’). Analysis of SGN number in sections through the mid-base region of the cochlea revealed no significant difference in young (2116.5+/-268.1, N=3, mean+/-standard error of the mean, SEM) *versus*. old (2064.3+/-158.4, N=3, mean+/-SEM) animals, assessed by staining for the pan-neuronal marker HuD. Likewise, double labeling for tdTomato revealed no significant changes in the proportion of Ib/Ic SGNs with age (Supplementary Fig. 1d, young=0.543+/-0.04, old=0.543+/-0.03, mean+/-SEM, p=0.65, one-sided Mann-Whitney). Collectively, these findings suggest that the observed synapse loss is not secondary to SGN cell death.

Quantitative analysis revealed that most surviving synapses were associated with Ia SGNs. To quantify the proportion of surviving synapses that belong to Ia (tdTomato-) versus Ib/Ic (tdTomato+) SGNs, we adapted an existing strategy ^25^ to identify synapses, defined by apposition of CtBP2 and Homer1 signal, and to assign SGN subtype identity, defined by assessing whether the Homer1+ punctum was associated with a tdTomato+ (Ib/Ic) or tdTomato-(Ia) terminal (analysis pipeline detailed in Supplementary Fig. 2, 3). A synaptic index (SI) was calculated by quantifying the total number of apposed pre-and post-synaptic puncta and dividing by the number of IHCs. Regardless of whether pre- or post-synaptic puncta were used to anchor the analysis, mean synaptic indices were significantly lower in old animals in comparison to young animals (Fig. 1e) (**ribbon-based SI:** young=18.1+/-0.8, old=11.8+/-0.9, mean+/-SEM, p=0.001, one-sided Mann-Whitney; **PSD-based SI:** young=17.3+/-0.9, old=11.3+/-0.8; mean+/-SEM, p=0.003, one-sided Mann-Whitney). Among the surviving synapses, the proportion belonging to Ib/Ic SGNs was significantly reduced in old (0.28+/-0.04) *versus*. young animals (0.45+/-0.03; p=0.005, one-sided Mann-Whitney) (Fig. 1f). Similar results were obtained by manual scoring and using a custom Python-based automated classification algorithm that assigns molecular identity based on tdTomato intensity within the reconstructed Homer1 surface (Supplementary Fig. 4). Collectively, these data show that Type Ia SGNs are the most resilient to age and that enhanced loss of Ib/Ic synapses precedes SGN cell death.

### Ib/Ic SGN synapses are disproportionally lost following noise exposure

To investigate whether Ib/Ic SGNs are also more vulnerable to noise-induced synaptopathy, we exposed *Ntng1*^*Cre*^*;Ai14* mice (8-9 weeks of age) to moderately intense sound stimuli, which cause permanently reduced auditory responses independent of hair cell damage (Fig. 2a). Responses to sound were evaluated by measuring ABRs, which reflect the collective activity of neurons upon presentation of stimuli of different frequencies and intensities^26^. The ABR threshold is the lowest intensity at which the peak in the first waveform (wave-I) is detected, and the amplitude of this response reflects the sum of SGN activity at that intensity. Increasing the sound intensity of the stimulus results in larger wave-I amplitudes as more SGNs are activated. Extremely intense stimuli can permanently elevate ABR thresholds, whereas more moderate stimuli cause elevations in ABR thresholds one day after exposure that return close to baseline after one week and are hence “temporary” threshold shifts. Wave-I amplitudes, however, do not fully recover because synapses are permanently lost^15^, predominantly in higher frequency regions of the cochlea. Thus, this Temporary Threshold Shift (TTS) paradigm is a useful way to investigate synaptopathy.

**Fig. 2:**
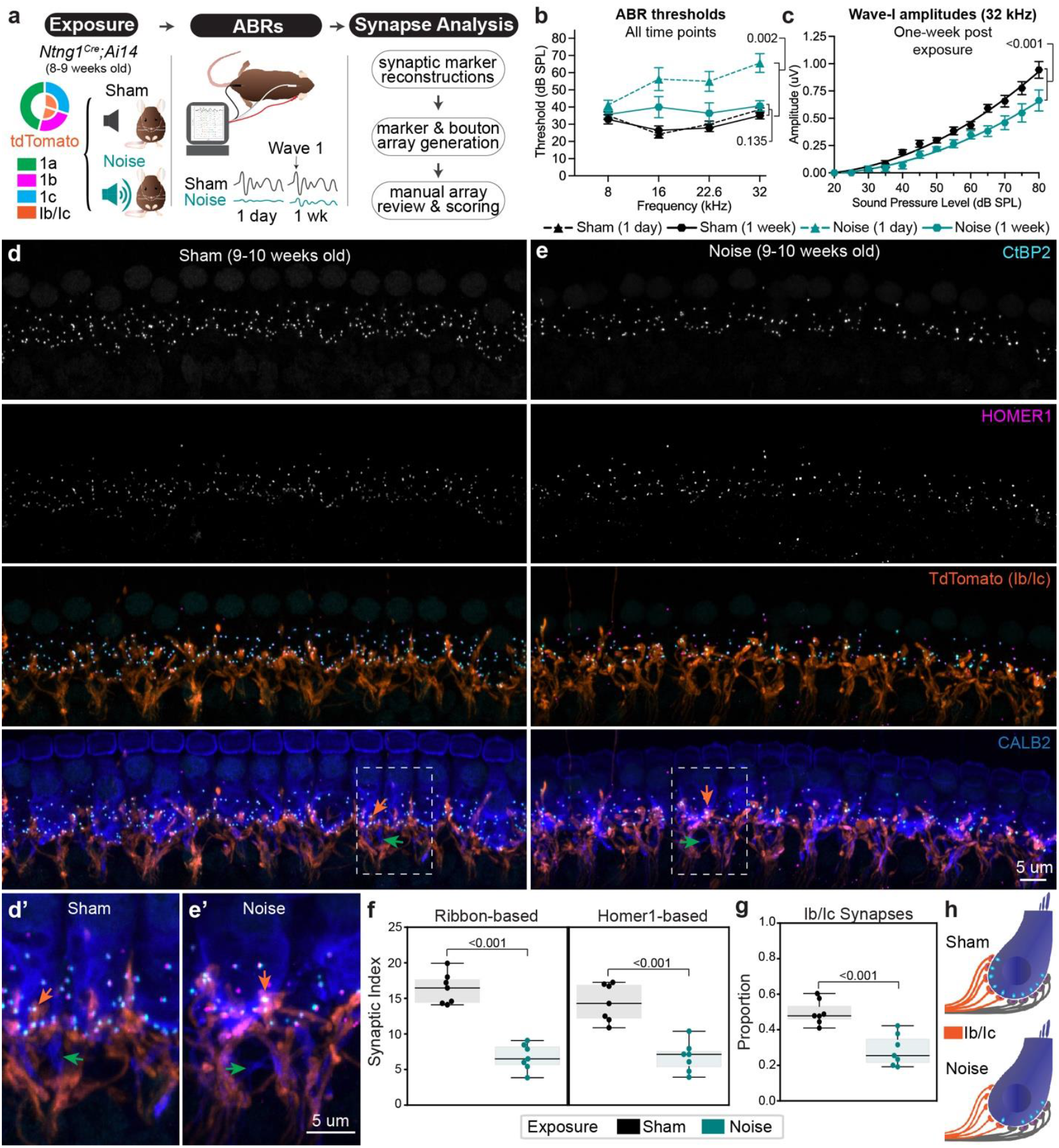
Noise-induced synaptopathy primarily affects Type Ib/Ic synapses. **a** Overview of experimental paradigm for inducing and analyzing synaptopathy with noise exposure in *Ntng1*^*Cre*^*;Ai14* reporter mice. **b** Thresholds for eliciting an ABR to a pure tone stimulus in sham and noise exposed mice at either one day or one week post exposure. Statistical comparisons show significantly elevated thresholds at the frequency analyzed for synapses (32 kHz) one day after but no significant difference one week after exposure (unpaired t-test). **c** Amplitudes of wave-I ABR responses at 32 kHz for each sound intensity tested. Curved line represents a second order polynomial, least squares fit of group means for each sound intensity tested. Statistical analysis is a comparison of non-linear fits. Individual dots in **b** and **c** represent group means (N=7 each) and whiskers represent SEM. **d, e** Representative high-power micrographs used for synapse and terminal analysis from sham and noise exposed mice, respectively. Dashed outlines correspond to regions magnified in **d’** and **e’** to highlight terminal morphology. **e** Quantification of synaptic density using either ribbon-based or Homer1-based approaches. **g** Proportion of Ib/Ic synapses as quantified from Homer1-based analysis. For **e** and **f**, each dot corresponds to a single field of view (one per animal), boxes represent interquartile ranges, horizontal lines represent the median, box height represents the interquartile range (IQR), and whiskers extend to data points farthest from the median but within 1.5x the IQR from the box edge. Statistical analyses performed with one-sided Mann-Whitney tests. **h** Schematized subtype-specific synapse loss following noise exposure extrapolated from quantification in **f** and **g**. Arrows for all panels: orange=Type Ib/Ic synapses, green=fibers with highest levels of CALB2.

We found that Type Ib/Ic synapses account for the vast majority of the total synapse loss in noise exposed *Ntng1*^*Cre*^*;Ai14* animals. Since genetic background influences resilience to noise exposure^27^, we first confirmed that *Ntng1*^*Cre*^*;Ai14* mice hear normally at baseline and reliably experience TTS after a two-hour exposure to 8-16 kHz octave band noise at 100 decibels sound pressure level (dB SPL) (Fig. 2b). Experimental mice (N=7) were compared to sham controls (N=7), which were handled identically and placed into the soundproof chamber except that no sound was played (Fig. 2a). ABR thresholds in noise exposed mice were significantly elevated relative to controls one day after exposure (**sham:** 38.57+/-5.46 dB SPL, **exposed:** 65.63+/-0.92 dB SPL, mean+/-SEM, p=0.002, unpaired t-test) but thresholds between the two groups were statistically indistinguishable at one week post-exposure (Fig. 2b) (**sham:** 35.00+/-1.89 dB SPL, **exposed:** 40.71+/-2.98 dB SPL, mean+/-SEM, p=0.135, unpaired t-test). Consistent with the presence of synaptopathy, wave-I amplitudes to pure tone stimuli of increasing sound intensity resulted in significantly different growth curves between noise and sham exposed mice (Fig. 2c) (R-squared sham=0.852, noise=0.762, mean+/-SEM, p<0.001, non-linear regression described in Methods).

Indeed, CtBP2+ and Homer1+ puncta were sparser in cochleae from noise exposed (Fig. 2e) versus sham exposed animals (Fig. 2d). Notably, the density of Type Ib/Ic terminals appeared relatively similar between the groups (Fig. 2d’ and 2e’), consistent with the persistence of SGN peripheral processes close to the IHC in electron microscopy (EM) reconstructions^28^. Quantification of the SI in the 32 kHz region using both ribbon- and PSD-based approaches confirmed that synaptic density was significantly reduced (Fig. 2f). On average, about half of the synapses were lost from noise exposed animals relative to sham controls (**ribbon-based SI:** sham=16.4+/-0.8 versus noise=6.7+/-0.7, mean+/-SEM, p<0.001; **Homer1-based SI:** sham=14.4+/-1.0 versus noise=6.8+/-0.8, p<0.001, one-sided Mann-Whitney). Ib/Ic synapses account for most of this loss, as the proportion of Ib/Ic (tdTomato+) synapses was significantly reduced in noise exposed mice (0.29+/-0.03) relative to sham controls (0.50+/-0.03) (mean+/-SEM, p<0.001, one-sided Mann-Whitney) (Fig. 2g). A small proportion of synapses in each group were excluded as they could not be definitively scored as being tdTomato+ (Supplementary Fig. 5a) (sham=0.09+/-0.01, noise=0.07+/-0.01, mean+/-SEM, p=0.228, one-sided Mann-Whitney). Assigning Ib/Ic status to these synapses did not alter the conclusions of the analysis (Supplementary Fig. 5b), further supported by an independent assessment based on tdTomato intensity (Supplementary Fig. 4). When placed in the context of a model IHC that starts with 10 synapses, this analysis predicts that four out of five Type Ib/Ic SGNs lose their synapses, whereas only one out of five Ia’s appears to be affected (Fig. 2h). Thus, Ia SGNs are more resilient than Ib/Ic SGNs to both noise-induced and age-related synaptopathy.

### Post-synaptic Homer1 volumes shift with age and noise in a subtype-specific manner

The identification of genetically defined subsets of vulnerable synapses offers an opportunity to systematically compare properties that might relate to their vulnerability. Previous analyses revealed a variety of changes in pre- and post-synaptic puncta volumes that differ along the pillar-modiolar axis and might therefore relate to SGN identity^25^. Because volume gradients are often subtle and can fluctuate depending on species^29^ and age^21^, subtype-specific differences in synaptic morphologies following noise exposure may be masked by anatomical variations. Additionally, it remains unclear whether vulnerable synapses show the same features after noise exposure as they do with age. To address these questions, we analyzed age- and noise-related changes in pre- and post-synaptic volumes in *Ntng1*^*Cre*^*;Ai14* mice.

Consistent with previous analyses along the pillar-modiolar axis, Type Ib/Ic Homer1 volumes were significantly smaller than those belonging to Type Ia SGNs at baseline and this difference persisted both with age and with noise exposure (**young:** Ib/Ic=1.08+/-0.02, Ia=1.22+/0.02, mean+/-SEM, p<0.001, two-sided Mann-Whitney; **old:** Ib/Ic=1.11+/-0.05, Ia=1.50+/-0.04, mean+/-SEM, p<0.001, two-sided Mann-Whitney; **sham:** Ib/Ic=1.08+/-0.02, Ia=1.27+/-0.03, p<0.001; **noise:** Ib/Ic=0.89+/-0.03 Ia=1.25+/-0.03, mean+/-SEM, two-sided Mann-Whitney). (Fig. 3a). Overall, post-synaptic volumes increased significantly with age (young=1.16+/-0.02, old=1.38+/-0.03, mean+/-SEM, p<0.001, two-sided Mann-Whitney) but not with noise (sham=1.16+/-0.02, noise=1.14+/-0.02, mean+/-SEM, p=0.969, two-sided Mann-Whitney) (Fig. 3b). However, Homer1 volumes changed differently depending both on SGN identity and the nature of sound exposure (Fig. 3c). For instance, Ia Homer1 volumes were significantly larger in old animals than in young ones (p<0.001, two-sided Mann-Whitney), whereas Ib/Ic Homer1 volumes did not increase significantly (p=0.207, two-sided Mann-Whitney). Unexpectedly, these trends were reversed in sham *versus*. noise exposed animals, where Type Ia volumes did not change significantly (p=0.441, two-sided Mann-Whitney) and Type Ib/Ic volumes became smaller (p<0.001, two-sided Mann-Whitney). These findings suggest that in the case of age, Ia synapses may build up their PSDs to make them more resilient, whereas in the case of noise exposure, Ib/Ic PSDs both start smaller and become smaller still, but average Ia PSD volumes do not change.

**Fig. 3:**
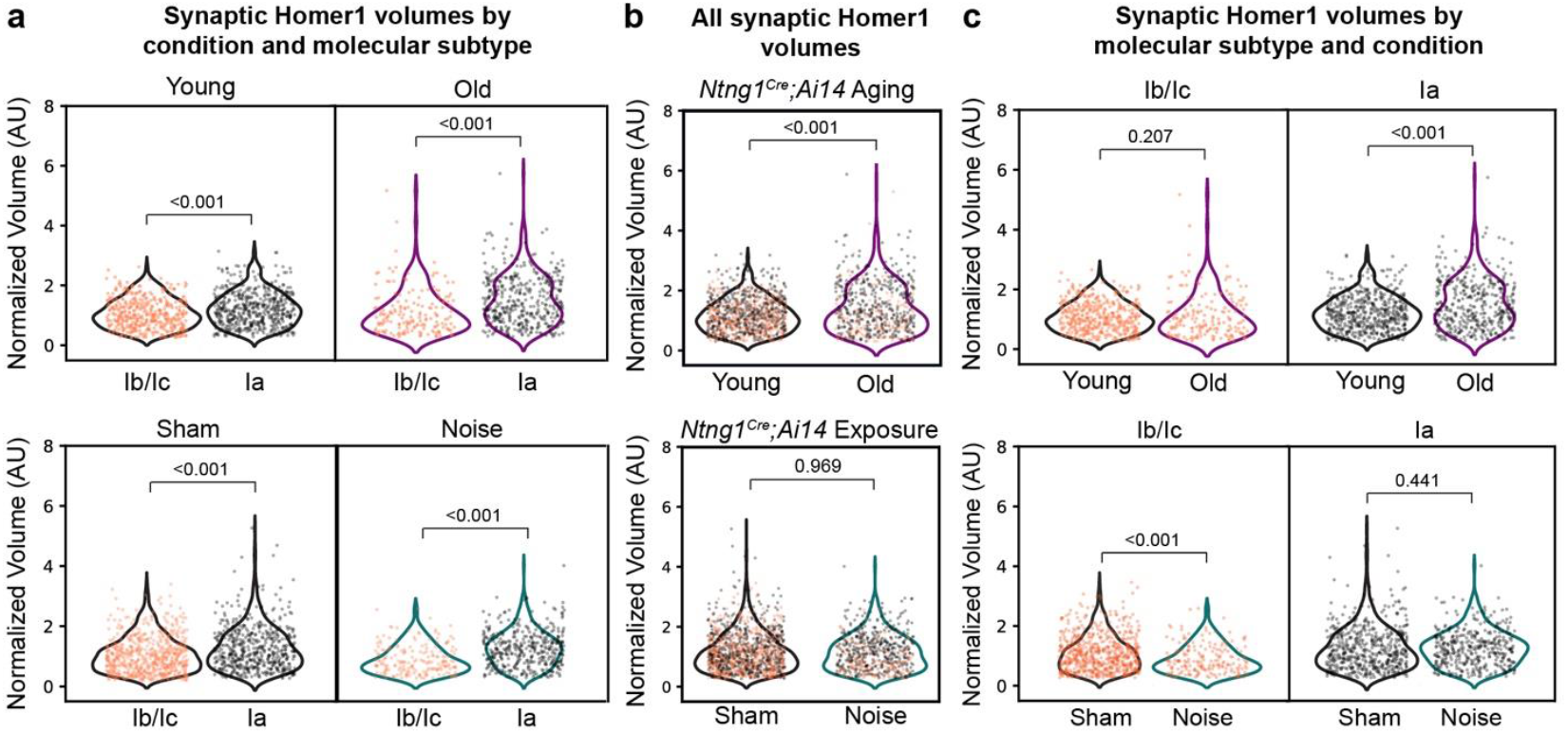
Subtype-specific changes in mean post-synaptic Homer1 volumes following age and noise. Violin plots of Homer1 volumes belonging to a synaptic pair, normalized to the median Homer1 volume of the control dataset for the group (one control volume per experiment, independent of molecular subtype). **a** Comparison of volumes between molecular subtypes (Ib/Ic versus Ia) within each condition (young versus old, top; sham versus noise, bottom). **b** All post-synaptic Homer1 volumes plotted by condition and color-coded by molecular subtype (orange = Ib/Ic, black = Ia). **c** Subtype-specific effects of aging (top) and noise (bottom) on Ib/Ic volumes (left column) and Ia volumes (right column). Two-sided Mann-Whitney test used for all statistical analyses.

### Density of Homer1 orphans increases with age and noise exposure

Noise-induced synaptopathy involves both an overall loss of ribbons that are apposed by PSDs and a transient increase in the number of ribbons and Homer1+ puncta that lack a partner (i.e. “ribbon orphans” and “Homer1 orphans”). Orphans occur at such low frequencies that it has been difficult to assess any subtype-specific changes rigorously. We observed orphans while scoring synaptic arrays and thus asked whether each orphan was associated with a tdTomato+ (i.e., Ib/Ic SGNs) or a tdTomato-terminal (i.e., Ia SGNs) (Fig. 4a, b). There were close to no ribbon orphans at baseline and only a modest increase with age or noise (young=0.02+/-0.02, old=0.12+/-0.07, mean+/-SEM, p=0.166, one-sided Mann-Whitney; sham=0.00+/-0.00, noise=0.48+/-0.11, mean+/-SEM, p<0.001, one-sided Mann-Whitney) (Fig. 4c). Homer1 orphans were more common in all conditions (young=0.98+/-0.24, old=1.61+/-0.19, mean+/-SEM, p=0.053, one-sided Mann-Whitney; sham=0.76+/-0.13, noise=2.17+/-0.37, mean+/-SEM, p=0.005, one-sided Mann-Whitney) (Fig. 4c). In the baseline condition, the vast majority of Homer1 orphans were within Type Ia terminals (Fig. 4d). However, with both age and noise exposure, the Type Ib/Ic Homer1 orphan proportion slightly, but significantly increased (young=0.07+/-0.03, old=0.24+/-0.06, mean+/-SEM, p=0.011, one-sided Mann-Whitney; sham=0.19+/-0.09, noise=0.37+/-0.03, mean+/-SEM, p=0.028, one-sided Mann-Whitney). Of note, the absolute number of PSD orphans remained low, averaging only one or two per IHC, so there were not enough to make robust comparisons of their volumes. However, a qualitative assessment suggests that Ia Homer1 orphan volumes are larger than the few that belong to Ib/Ic’s at baseline and this difference persists with age and noise exposure (Fig. 4e). Collectively these data suggest that the synapses made by SGN subtypes are differentially affected by age and noise (Fig. 4f). At baseline and with age, essentially all of the Homer1 orphans belong to Ia’s. However, with noise exposure, the new orphans that appear are most likely to come from Ib and/or Ic SGNs.

**Fig. 4:**
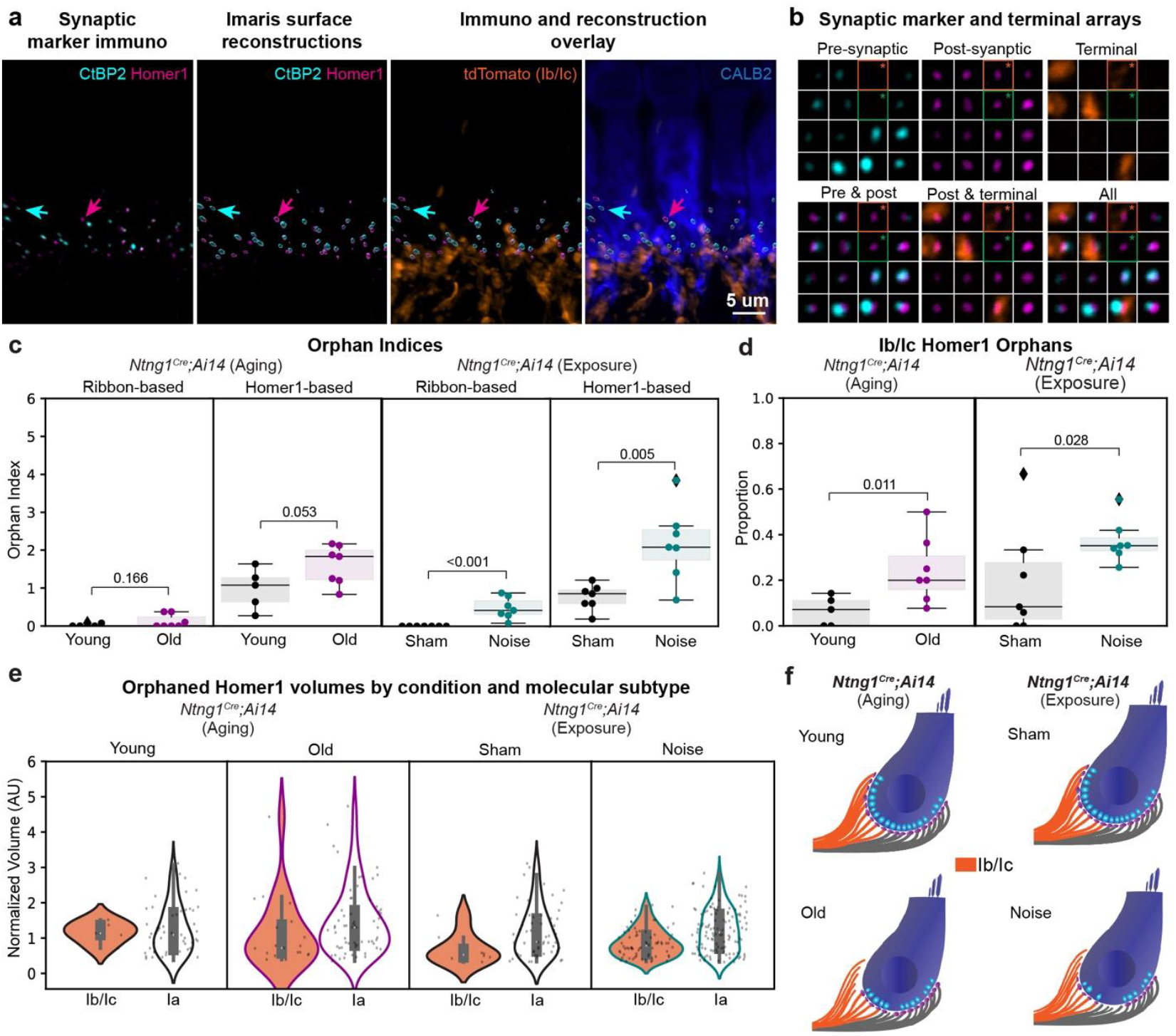
Molecular subtypes have different proportions of Homer1 puncta lacking presynaptic partners. **a** Imaris generated three-dimensional renderings from high magnification micrograph used for synapse and terminal analysis. First panel shows the fluorescent channels for the pre-synaptic marker CtBP2 and post-synaptic marker Homer1. Second panel shows the surface reconstructions generated for each channel after removal of surfaces that are outside of the IHC boundary. Third and fourth panels show these surfaces overlaid with the actual fluorescence channels for tdTomato (Ib/Ic’s) and CALB2 (IHCs, Ia’s). Cyan arrows point to a CtBP2 puncta lacking a Homer1 partner (i.e., “ribbon orphan”). Magenta arrows point to a Homer1 puncta lacking a CtBP2 puncta (i.e., “Homer1 orphan”). **b** Representative synaptic and terminal arrays highlighting two Homer1 orphans, one belonging to a Ib/Ic terminal (orange box and asterisk) and one belonging to a Ia terminal (green box and asterisk). Each thumbnail image is 1.5 um x 1.5 um. **c** Comparison of ribbon orphan and Homer1 orphan densities across ages and noise exposure, respectively. **d** Comparisons of Type Ib/Ic Homer1 orphan proportion across age and noise exposure, respectively. For **c** and **d** each data point represents a single field of view (one field of view per animal), boxes represent interquartile ranges (IQRs), horizontal lines represent the median, box height represents the IQR, and whiskers extend to data points farthest from the median but within 1.5x the IQR from the box edge. Diamonds mark data points outside whiskers classified as potential outliers by Python Seaborn package. **e** Comparison of control-median normalized orphaned Homer1 volumes between molecular subtypes (Ib/Ic versus Ia) within each condition (i.e., young, old, sham exposed, noise exposed). **f** Model of synapse loss and increased orphan density with age and noise as extrapolated from indices and proportions shown in **c** and **d**. Statistical analyses performed with one-sided Mann-Whitney tests for **c** and **d**, including data points classified as potential outliers. Statistical analyses are not possible for **e** due to limited and/or uneven sample sizes.

### Complementary patterns of resilience and vulnerability in Type Ia versus Ic SGNs

Given the clear differences in vulnerability revealed by analysis of *Ntng1*^*Cre*^*;Ai14* animals, we sought to make a more direct comparison between the most (Ic) and least (Ia) vulnerable neurons. We therefore repeated the TTS-inducing noise exposure paradigm using tamoxifen-inducible Cre reporter lines that drive tdTomato expression in either Ia and Ib (*Calb2*^*CreERT2*^) or just Ic (*Lypd1*^*CreERT2*^) SGNs^11^. Mice from both reporter lines were injected with a dose of tamoxifen previously shown to label mostly Ia’s and only a few Ib’s in *Calb2*^*CreERT2*^*;Ai14* and primarily Ic’s in *Lypd1*^*CreERT2;*^*Ai14* ^11^. Tamoxifen was delivered to mice at P26-P30 (*Lypd1*^*CreERT2;*^*Ai14* N=12, *Calb*^*CreERT2;*^*Ai14* N=16), four weeks prior to noise exposure. ABRs were conducted at one- and seven-days post-noise exposure, as for *Ntng1*^*Cre*^*;Ai14* experiments (Fig. 5a). Both lines exhibited TTS, with thresholds returning close to baseline one week after exposure, with minor differences in the extent of recovery (Supplementary Table 1). Ia terminals were noticeably larger than the Ic terminals (Fig. 5b, white arrows), providing additional anatomical evidence of their identities, since high SR axons have greater calibers than low SR axons^30^. TdTomato+ terminals in the *Calb2*^*CreERT2*^*;Ai14* mice frequently overlapped with CALB2 and were primarily on the pillar side of the IHCs (Fig. 5c), but as with prior reports, this innervation pattern was not strictly observed^11^. By contrast, tdTomato+ terminals in *Lypd1*^*CreERT2*^*;Ai14* mice did not overlap with CALB2 and were primarily on the modiolar side, consistent with a Ic identity. In both lines, a subset of Homer1+ puncta were localized within tdTomato+ terminals and apposed by CtBP2+ ribbons. Only a subset of Ia SGNs were labeled in *Calb*^*CreERT2;*^*Ai14* mice, whereas coverage of Ic SGNs was more complete in *Lypd1*^*CreERT2*^*;Ai14* mice.

**Fig. 5:**
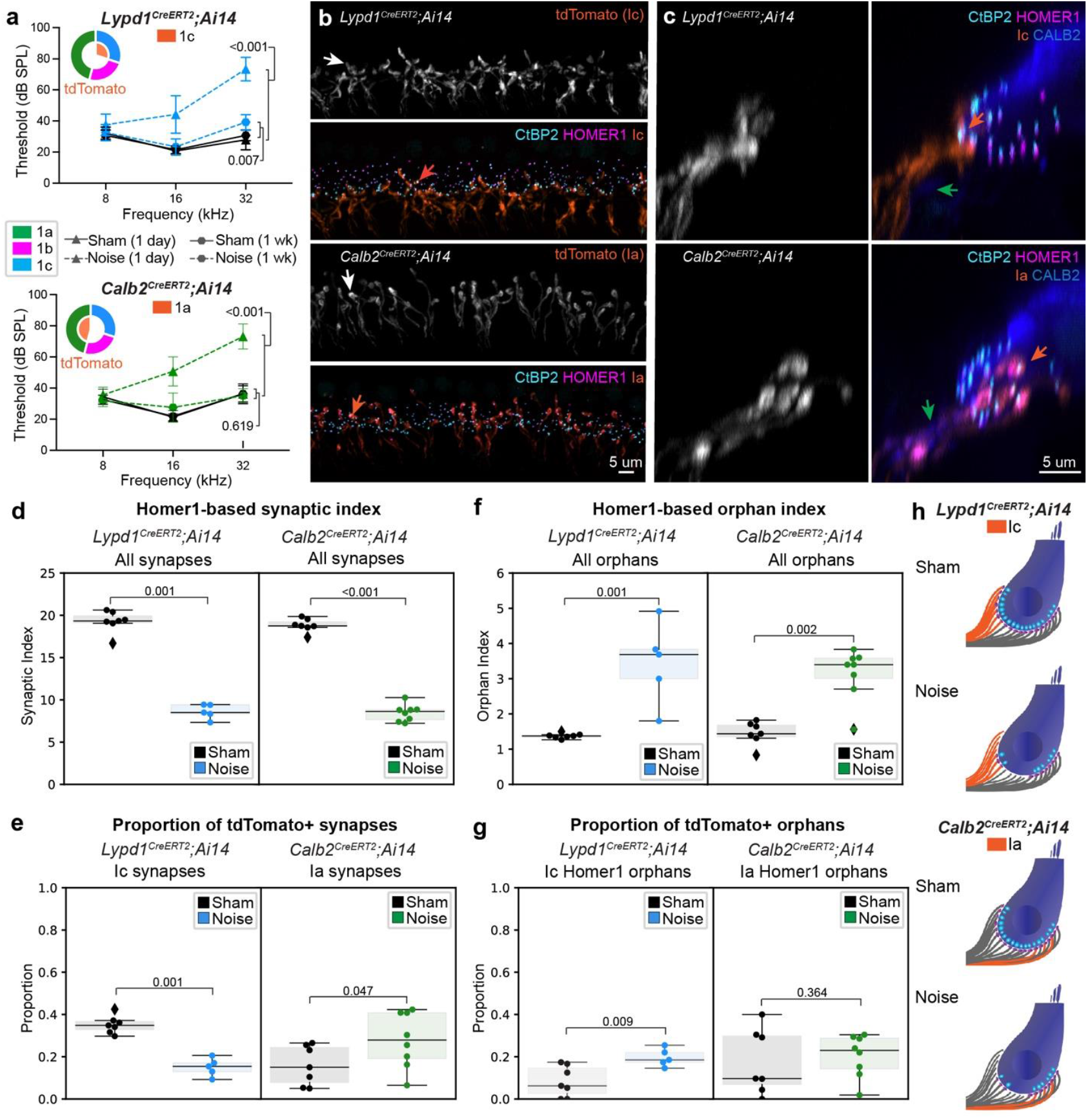
Resilience of Type Ia SGNs to noise-induced synaptopathy. **a** Thresholds for eliciting an ABR to a pure tone stimulus in sham and noise exposed mice from Ic and Ia reporter lines (*Lypd1*^*CreERT2*^*;Ai14* and *Calb2*^*CreERT2*^*;Ai14*, respectively) at either one day or one week post exposure. Statistical comparisons between sham and exposed animals at one day and one week post exposure at 32 kHz (i.e., the tonotopic region analyzed for synaptic density) (unpaired t-test for all comparisons). **b** Representative high-power micrographs from each reporter line used for synapse and terminal analysis. Blue and green arrows highlight terminal morphology in each reporter line. **c** Maximum-intensity projections of virtually sliced z-stacks from each line (10 um wide section) shows tdTomato terminals in Ic reporter line primarily innervate the modiolar region of the inner hair cell (IHC) whereas those terminals primarily innervate the pillar region in the Ia reporter line. In both cases, tdTomato+ terminals overlap with anatomically distinct subsets of Homer1 puncta (i.e., those on the modiolar versus pillar side). For **b** and **c** arrows: white=terminals, orange=tdTomato+ synapses, green=fibers with highest levels of CALB2. **d** Comparison of Homer1-based synaptic indices in sham versus noise exposed mice, for each reporter line. **e** Reporter-line specific differences in tdTomato+ synapse proportion. **f** Density of Homer1 orphans in sham versus noise exposed mice for each reporter line. **h** Comparison of Homer1 orphan density within each reporter line with noise exposure. For **d**-**h**, each data point represents a single field of view (one field of view per animal, Ic N=7 sham, 5 noise; Ia N=7 sham, 8 noise), boxes represent interquartile ranges, horizontal lines represent the median, box height represents the interquartile range (IQR), and whiskers extend to data points farthest from the median but within 1.5x the IQR from the box edge, diamonds mark data points outside whiskers and are classified as potential outliers by Python Seaborn package; one-sided Mann-Whitney test used for all statistics and include data points classified as potential outliers. **f** Schematized synapse loss and increase in orphan density for each reporter line and molecular subtype.

Upon exposure to noise, both lines experienced significant synapse loss that was primarily restricted to the Ic population. Noise exposure reduced the SI to similar extents in the 32 kHz region in both reporter lines (***Lypd1***^***CreERT2;***^***Ai14* mean SI:** sham=19.26+/-0.48, noise=8.60+/-0.39, p=0.001; ***Calb2***^***CreERT2;***^***Ai14* mean SI:** sham=18.83+/-0.30, noise=8.45+/-0.35, mean+/-SEM, p<0.001, one-sided Mann-Whitney) (Fig. 5d). In comparison to sham controls, noise exposed *Lypd1*^*CreERT2;*^*Ai14* mice had a significantly smaller proportion of synapses made by tdTomato+ terminals, which come from Ic SGNs in this line (**mean Ic synapse proportion:** sham=0.35+/-0.02, noise=0.15+/-0.02, mean+/-SEM, p=0.001, one-sided Mann-Whitney) (Fig. 5e). By contrast, in noise exposed *Calb2*^*CreERT2*^*;Ai14* mice, a significantly greater proportion of surviving synapses were associated with tdTomato+ terminals, which come from mostly Ia SGNs in this line (**mean Ia synapse proportion:** sham=0.16+/-0.04, noise=0.28+/-0.05, mean+/-SEM, p=0.047, one-sided Mann-Whitney) (Fig. 5e). Thus, these experiments directly demonstrate the resilience of Type Ia SGNs to noise-induced synaptopathy. Similar results were obtained when synapses with uncertain tdTomato status were included (Supplementary Fig. 5) and by automated analysis of tdTomato+ intensities (Supplementary Fig. 4).

Analysis of Homer1 orphans reproduced what was observed in *Ntng1*^*Cre*^*;Ai14* mice. In both reporter lines, the mean orphan index was significantly greater in noise exposed animals relative to sham controls (***Lypd1***^***CreERT2;***^***Ai14*:** sham=1.27+/-0.03, noise=3.45+/-0.51, mean+/-SEM, p=0.001, one-sided Mann-Whitney; ***Calb2***^***CreERT2;***^***Ai14*:** sham=1.45+/-0.12, noise=3.15+/-0.26, mean+/-SEM, p=0.002, one-sided Mann-Whitney) (Fig. 5f). This increase largely reflects the addition of Type Ic Homer1 orphans, since the proportion of tdTomato+/Ic Homer1+ orphans is significantly higher in noise exposed *Lypd1*^*CreERT2;*^*Ai14* animals relative to sham controls (sham=0.08+/-0.03, noise=0.20+/-0.02, mean+/-SEM, p=0.009, one-sided Mann-Whitney) but the proportion of tdTomato+/Ia synapses does not change in *Calb2*^*CreERT2;*^*Ai14* animals (sham=0.18+/-0.05, noise=0.20+/-0.04, mean+/-SEM,p=0.364, one-sided Mann-Whitney) (Fig. 5g). Consistent with what was observed for *Ntng1*^*Cre*^*;Ai14*, these experiments confirm that Ic SGNs are more likely to lose synapses and hence retain orphans than Ia SGNs following noise exposure (Fig. 5h).

### Shifting Type Ib/Ic SGNs towards a Ia identity protects synapses from noise

The relative resilience of Ia synapses to noise exposure suggests that intrinsic molecular differences may be responsible. Alternatively, there may be something about the nature of the synapses in the pillar region of the IHC that makes them less susceptible to glutamate-induced excitotoxic damage^31^. For example, lateral olivocochlear efferent neurons provide more innervation to afferents positioned on the modiolar side than on the pillar side and de-efferentation leads to changes in ribbon and glutamate receptor patch sizes^32^.

To distinguish among these possibilities, we altered the proportions of Type I SGN subtypes by conditionally deleting *Runx1* from SGNs using *Bhlhe22*^*Cre*^ (aka *bhlhb5*^*Cre*^) mice, which mediates recombination in SGNs but not in other cell types in the cochlea^33^. *Runx1* encodes a transcription factor that promotes Ib and Ic SGN fates^34^ and its conditional deletion from SGNs increases the proportion of Ia-like SGNs at the expense of Ib and Ic SGNs, as assessed by single cell RNA-sequencing^34^. Importantly, total SGN density did not change and SGN terminals still filled the basolateral surface of the IHC. However, gradients of ribbon and PSD size changed, such that terminals on the modiolar side of the IHC took on Ia-like properties, offering an opportunity to assess their resilience to noise.

Analysis of synapse loss in noise exposed *Runx1* conditional knock-out (CKO, *Bhlhe22*^*Cre/+*^*;Runx1*^*flox/flox*^) and control (CON, *Bhlhe22*^*+/+*^*;Runx1*^*flox/flox*^) animals demonstrated that molecular differences among SGN subtypes impact vulnerability. Mice of this genetic background were more sensitive to auditory damage and experienced permanent threshold shifts even with a 97.5 dB SPL stimulus, with similar effects in both genotypes (**CON noise:** N=10, mean threshold=83.64+/-0.70, **CKO noise:** N=9, mean threshold=81.88+/-3.13, mean+/-SEM, p=0.065, mixed-effects analysis with multiple comparisons) (Fig. 6a, b). As IHC-localized CtBP2+ puncta are an excellent proxy for synaptic index and are rarely found as orphans (Fig. 4 and from others^35,36^), we streamlined the analysis by quantifying only pre-synaptic ribbons. We found that ribbon density was diminished following noise exposure in both genotypes. However, CKO mice retained significantly more ribbons than their CON littermates following noise exposure (**mean ribbon index, CON noise:** N=10, mean=8.50+/-0.68, **CKO noise:** N=9, mean=11.66+/-1.00, mean+/-SEM, p=0.023, one-sided Mann-Whitney) (Fig. 6c,d). Quantification of the SI in a subset of preparations similarly showed more synapses in CKO relative to CON noise exposed animals (**CON noise:** N=8, mean=9.09+/-0.70, **CKO noise:** N=5, mean=12.29+/-1.23, mean+/-SEM, p=0.047, one-sided Mann-Whitney).

**Fig. 6:**
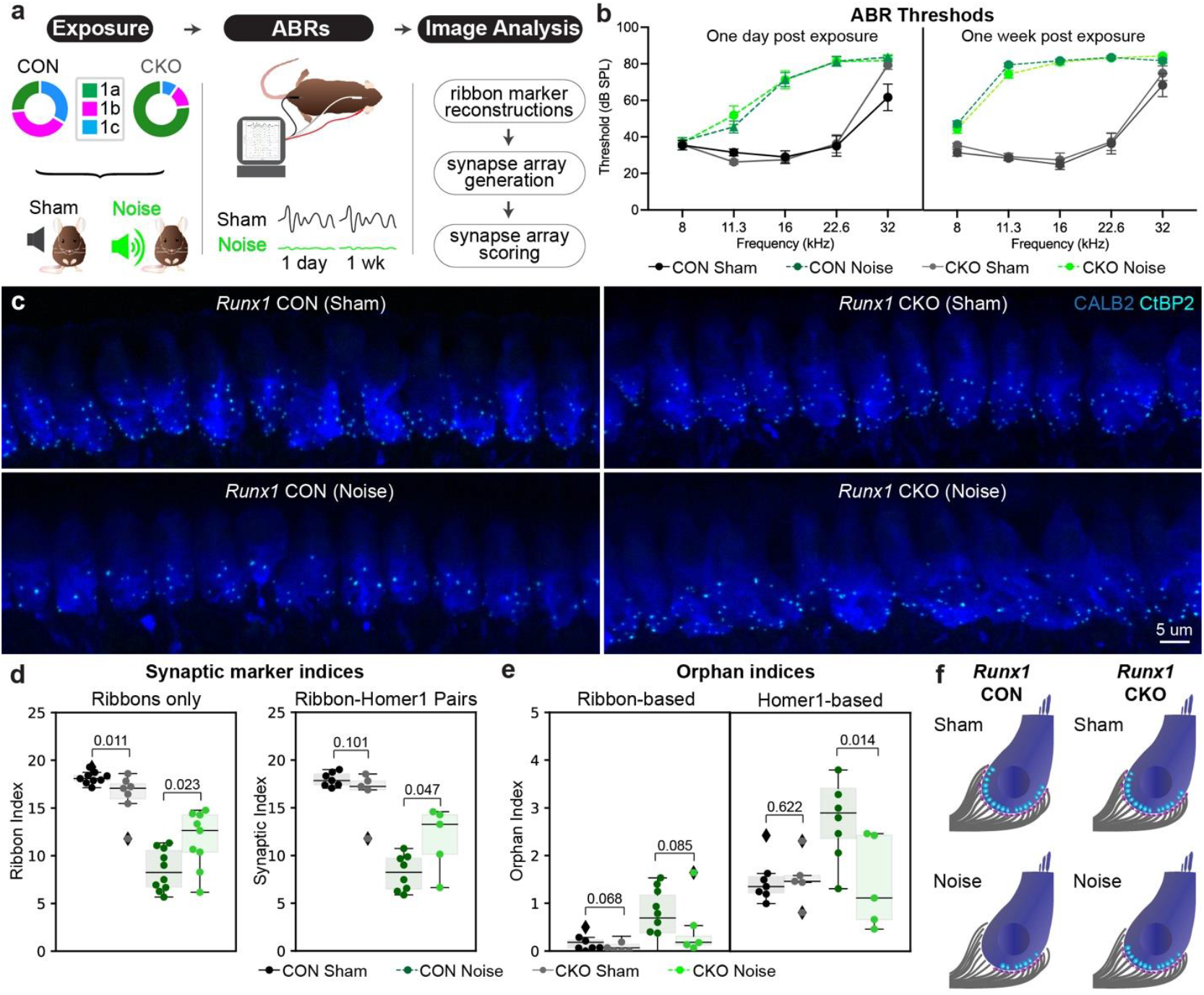
Conditional deletion of *Runx1* protects against noise-induced synapse loss. **a** Overview of experimental paradigm for inducing synaptopathy using moderate noise exposure in *Runx1* conditional knock-out (CKO) and control (CON) mice. **b** Thresholds for eliciting an ABR to a pure tone stimulus in sham and noise exposed mice (CON and CKO) one day and one week after exposure. **c** Representative high-power micrographs used for ribbon analysis in sham and noise exposed animals of both genotypes. **d** Quantification of the density of ribbons for each condition (ribbon index, left) and synapses (right) in the subset of animals for which Homer1 immunolabelling was possible. **e** Quantification of CtBP2 orphans (ribbon-based) and Homer1 orphans. For **d** and **e**, each data point represents a single field of view (one field of view per animal, ribbon-only analysis CON N=9 sham, 7 noise, CKO N=7 sham, 9 noise; ribbon and Homer1 analysis CON N=7 sham, 5 noise, CKO N=5 sham, 5 noise), boxes represent interquartile ranges, horizontal lines represent the median, box height represents the interquartile range (IQR), and whiskers extend to data points farthest from the median but within 1.5x the IQR from the box edge, diamonds mark data points outside whiskers and are classified as potential outliers by Python Seaborn package; one-sided Mann-Whitney test used for all statistics to compare effects of genotype within an experimental condition and include data points classified as potential outliers. **f** Model of synapse loss and changes in the number of Homer1 orphans as they occur with noise exposure across the genotypes, as extrapolated from the results plotted in **d** and **e**.

Analysis of orphaned synaptic markers further supported the conclusion that the shift towards a Ia identity made synapses more resilient. Although the number of orphans after noise exposure increased in both genotypes, CKO mice acquired fewer ribbon orphans (**mean orphan index, CON:** noise=0.91+/0.16, **CKO:** noise=0.51+/-0.29, mean+/-SEM, p=0.085, one-sided Mann-Whitney) and fewer Homer1 orphans (**mean Homer1 orphan index, CON:** noise=3.12+/-0.52, **CKO:** noise=1.43+/-0.43, mean+/-SEM, p=0.014, one-sided Mann-Whitney) than CON mice (Fig. 6e). Of note, CKO mice start with slightly fewer synapses than their CON littermates but have the same baseline number of PSD orphans. These data suggest that following noise exposure, CON mice lose about 53% of their synapses and gain a significant number of Homer1 orphans (Fig. 6f). Only about 33% of synapses are lost in the CKO mice, which acquire essentially no new Homer1 orphans. This finding strongly supports the conclusion that gene expression differences between Ia and Ic SGNs are responsible for selective vulnerability to noise exposure.

## Discussion

Though it has long been hypothesized that low threshold, high SR SGNs are more resilient than high threshold, low SR SGNs, definitive proof was missing due to the dearth of specific molecular markers for the Type I SGN subtypes. Here, we used genetic approaches to demonstrate that Type Ia SGNs, which match the anatomical properties of low threshold, high SR SGNs, show a resilience to noise-induced and age-related synaptopathy that is not present in Ic SGNs, which match the anatomical properties of high threshold, low SR SGNs. Across three different Cre reporter lines that experienced the same degree of noise-induced synapse loss, the proportion of Ic synapses decreased and the proportion of Ia synapses increased. This was true regardless of whether Ia or Ic SGNs expressed tdTomato. Ia resilience reflects, at least in part, intrinsic molecular differences, as we also observed greater protection from cochlear synaptopathy when SGN identities and synaptic properties became more Ia-like. This fits with the discovery that noise exposure activates an ATF3/ATF4 cellular stress response pathway in Ia’s and not in Ib’s or Ic’s^37^. Ib SGN responses remain unclear, but our data suggest they are also more resilient than Ic’s, since more tdTomato+ synapses survived in *Ntng1*^*Cre*^*;Ai14* animals, where both Ib and Ic terminals are labeled, than in *Lypd1*^*CreERT2*^*;Ai14* animals, where only Ic terminals are labeled. Additionally, Ib’s are more likely to survive with age than Ic SGNs^8^. An important area for future investigation is identifying the cause of Ic vulnerability. We also observed enhanced loss of Ib/Ic synapses in old mice and this preceded any evidence of SGN cell body loss. Analysis of Homer1 orphans suggests that Type Ia synapses experience either less complete synapse loss or more efficient repair. Together, these findings suggest that age-related and noise-induced cochlear synaptopathy could be mitigated by specifically inducing Ia-like protective mechanisms in Ic SGNs.

Despite the well-established anatomical and physiological heterogeneity of IHC-SGN synapses, it has been challenging to uncover which molecular differences might influence their resilience^6,31,38^. In fact, although the gradients of ribbon and PSD volumes map onto the same axis as the Ia, Ib, and Ic terminals, the distributions of volumes are wide and there is considerable overlap. Further, the degree of heterogeneity varies across genetic backgrounds and can change depending on how the pillar-modiolar axis is defined^39^. Synaptic features also change after depletion of olivocochlear efferent innervation, highlighting the dynamic nature of the IHC-SGN synapse^32^. However, at least some of this heterogeneity appears to be intrinsically determined, as ribbon volume correlates with SGN identity^8^ and mean ribbon and PSD volumes shift predictably with SGN identity^34^ and when subtype-appropriate programs of gene expression are altered^40^. Consistent with these findings, we found that Homer1+ PSD volumes are largest in Ia SGNs and that Ia PSDs may be more resilient, though it is unclear whether the increased volumes play any role in this resilience.

Although the difference between Ia and Ic volumes increased with both aging and noise, closer analysis showed that the effects on the PSDs differ, raising the possibility that there are different molecular mechanisms for resilience and vulnerability. In the case of aging, the mean Ia Homer1 volume increased. By contrast, with noise, the mean Ib/Ic volume decreased. The noise-induced decrease in Homer1 volumes could be caused by either a complete loss of larger synapses or a uniform shrinking of PSDs that results in the complete loss of the smallest synapses. This change in volume could occur through the same pathways that normally maintain pools of glutamate receptors (AMPARs) in an activity-dependent manner^41^. Similarly, Type Ic SGN PSDs could shrink as glutamate receptors are endocytosed and go unreplaced. In support of this idea, AMPAR cycling begins within minutes of excessive noise exposure^42^, possibly as a synapse-protective response^35^. Hence, Type Ic SGNs might experience more receptor endocytosis under excitotoxic conditions, such that the damage becomes too severe to repair. Alternatively, Ia SGNs may be more efficient in rebuilding synapses from the orphans that remain. There is strong evidence for some degree of synapse repair^36^. For instance, EM reconstructions indicate that three out of five terminals lose their entire post-synaptic density at 24 hours post-exposure, but one out of the three that are lost will be rebuilt by one week post-exposure^28^.This may also explain the counterintuitive finding that Ia SGNs were more likely to be associated with PSD orphans both before and after exposure. The number of PSD orphans also increased in Ib/Ic SGNs but accounted for a much smaller proportion of all lost synapses. Perhaps Ia SGNs are more likely to retain a PSD and hence eventually repair the synapse.

The increase in Ia SGN Homer1 volumes may reflect a different type of protective response. Surviving SGN terminals in aged animals appear larger^21^ and, given our findings, most likely belong to Ia SGNs. Since our data revealed essentially no loss of Ia synapses with age, the increase in mean volume is likely due to an increase in the size of the PSD in surviving terminals, rather than a loss of small PSDs. Such a compensatory response occurs widely in the nervous system, where neurons frequently scale their PSDs in response to changes in pre-synaptic activity^41^. A possible role for similar scaling in the cochlea is supported by the fact that AMPAR patches are larger in mice that cannot release glutamate^36^. Age-related changes in auditory function are not limited to SGNs and multiple sources of damage could result in age-related reductions in glutamate release from IHCs. Thus, the observed increase in Type Ia Homer1 volumes could be a result of decreased glutamate release at the ribbon synapse that stems from age-related changes in cochlear function. Why such a mechanism is not evident in Ic synapses is unclear.

Another key contributor to variable IHC-SGN synapse resilience may be subtype-specific differences in calcium buffering properties. The current model of noise-induced synaptopathy is that excessive glutamate leads to calcium-induced excitotoxicity that overloads post-synaptic calcium buffering capacity^4,5^, perhaps pushing some synapses past the point where they can be repaired. Differences in the axon caliber and mitochondrial content of pillar versus modiolar terminals support the idea that Type Ib/Ic SGNs are more vulnerable to calcium dysregulation^28,30^. Mitochondria act as calcium sinks during synaptic transmission^43–45^, so more mitochondria could in theory result in greater calcium buffering capacity. Similarly, the majority of remaining terminals in aged mice were pillar-innervating and had larger mitochondrial volumes^21^, consistent with a Ia identity. More work is needed to determine how different subtypes handle calcium, how the number of mitochondria influences this response, and whether there are also subtype-specific differences in mitochondrial biology that cause functional differences in calcium buffering capacity and/or ATP production.

The identification of intrinsically different vulnerabilities among SGN subtypes provides a valuable entry point for therapeutic studies. The loss of Ic and perhaps also Ib synapses leads to clear differences in auditory responses that fit well with what we know about subtype-related differences in physiology. Low SR SGNs resist saturation at higher sound intensities and could be important for sound encoding in loud environments^46^. Although cochlear synaptopathy also affects humans^47^, there is not yet a proven human perceptual correlate. This may be a limitation of diagnostic power rather than a true fact of the biology, as patients with normal ABR thresholds can experience “hidden hearing loss,” a phenomenon in which hearing performance is poor despite normal abilities to detect sound^17,48^. This has been hypothesized to be due to a loss of Type Ib and/or Ic SGNs and our data in mice supports this. In these experiments, where ABR thresholds were normal, 32 kHz wave-I amplitudes at 70 dB SPL (the sound intensity of loud conversations) were significantly reduced by 30% and correlated with a 42% reduction in Type Ib/Ic synapses. What this degree of loss means for hearing function in humans and whether such insights could be harnessed to improve cochlear implant functionality is an important application of future therapeutics research. *Runx1* CKO mice confirmed that synaptic vulnerability can be tuned towards resilience by altering neuronal identity. This raises the possibility that certain genes or drugs might be able to reproduce Ia-related resilience and thus improve the ability to hear across a wide dynamic range, even when the environment is noisy. Additionally, molecular differences in the vulnerability or resilience of SGN synapses may lead to the identification of pathways that are relevant to other forms of synaptopathy, including Alzheimer’s Disease.

Our conclusions are supported by a multi-pronged genetic approach, but some limitations should be noted. First, our analysis was restricted to the 32 kHz region of the cochlea, and it is possible that the degree of the effect is different at other tonotopic positions, especially as there are slight differences in the proportion of Ia, Ib, and Ic SGN subtypes along the apical-basal axis. Another limitation is that we only tracked terminals that were filled with tdTomato in each reporter line and thus could only infer what happened to unlabeled terminals. This limited our ability to make strong conclusions about what happens to Ib synapses in particular. Additionally, some of the reductions in the Ib/Ic synapse density could be due to downregulation of the *Rosa26* locus or altered trafficking of tdTomato between the soma and terminal. For instance, if insufficient tdTomato was trafficked to the repaired terminal it could appear tdTomato-. With these potential confounds in mind, we designed our analysis to score terminals based on signal distribution rather than simply considering intensity. This approach appears to be robust, as our automated classification of subtype synapse proportions independently supported the conclusions from manual scoring. However, it remains possible that some terminals may have been misidentified because of the nature of the tdTomato signal. Additionally, although the proportion of HuD+ and tdTomato+ SGN somata in *Ntng1*^*Cre*^*;Ai14* animals did not decrease with age, the complex morphology of the ganglion makes it difficult to measure SGN density reliably in cochlear sections. Thus, while our data strongly suggest that there is a substantial degree of age-related synapse loss that cannot be explained entirely by the loss of SGNs, it is possible that some of the synapses are missing because the SGN itself is gone, not because of synaptopathy *per se*. Finally, there are several cell-extrinsic sources of protection that were not studied here, including neuroimmune response and efferent activity, which may make important contributions to the measured vulnerability^49,50^.

## Materials and Methods

### Mice

Animal work was conducted in compliance with protocols approved by the Institutional Animal Care and Use Committee at Harvard Medical School. Mice were housed in the Harvard Center for Comparative Medicine in groups of up to five animals and maintained on a 12hr light/dark cycle. All experiments used both male and female mice. Food and water were available *ad libitum. Cre* and *CreERT2* lines were maintained on mixed backgrounds crossed to CBA/CaJ (Jax strain 000654) to ensure heterozygosity for the age-related *Cdh23*^*753G>A*^ mutation (i.e., *Cdh23*^*753G/753A*^)^51^. The following previously published mouse lines were used: *Netring1*^*Cre*^ (MGI: 6740547) ^52^; *Rosa26*^*LSL-tdTomato*^ (Jax: 007914)^53^; *Bhlhe22*^*Cre*^ (MGI:4440745)^54^; *Runx1*^*flox*^ (MGI:4358510). *Ntng1*^*Cre*^*;Ai14* mice used for aging studies (experimental and controls) were maintained as homozygous for the age-related *Cdh23*^*753G>A*^ mutation (i.e., *Cdh23*^*753A/753A*^). *CreERT2* activity in *Calb2*^*CreERT2*^*;Rosa26*^*LSL-tdTomato*^ and *Lypd1*^*CreERT2*^*;Rosa26*^*LSL-tdTomato*^ mice was induced with a single intraperitoneal injection of tamoxifen (0.1 mg/g body weight) between P26-P31. Tamoxifen (Sigma Aldrich T5648-1G) was dissolved in corn oil to a final concentration of 20mg/mL.

### Noise Exposure

Noise exposure was administered to animals in a custom plexiglass trapezoidal box located inside a tabletop sound-attenuated chamber. Acoustic stimuli were delivered from a speaker at the top of the chamber. To ensure even sound exposure, animals were placed in individual wire mesh boxes on a mesh platform located in the center of the chamber. Sound pressure levels were measured with a 1/4” free-field microphone (PCB 378C01) which was calibrated prior to each exposure session (Larson-Davis CAL200). Stimuli consisted of an 8–16 kHz octave-band noise for 120 minutes. Up to 4 animals were exposed at a time with the light on inside of the box. Sham exposed animals were placed in the wire mesh boxes within the plexiglass chamber for the same duration as noise exposed animals. Because noise/sham exposures were administered across multiple days, special care was taken to mix animals of different genotypes and sex for each exposure when possible. Sound intensity was recorded for the duration of the exposure to ensure constant noise level.

### Cochlear Function Testing

Mice were anesthetized with an initial intraperitoneal injection of ketamine (100 mg/kg) and xylazine (10 mg/kg). Meloxicam (1 mg/kg) was administered intraperitoneally for analgesia prior to assessments. Animals were placed on a 37°C heating pad (ATC1000, World Precision Instruments) and additional ketamine (30-40 mg/kg) was administered as needed to maintain the anesthetic plane throughout the procedure, except in the case of *Ntng1*^*Cre*^*;Ai14* experiments. *Ntng1*^*Cre*^*;Ai14* mice exhibit a heightened sensitivity to anesthesia such that subsequent ketamine administration often leads to death. Therefore, to preserve group sizes for those experiments, if animals woke up during a measurement, the session was terminated, and the mouse was placed on a heating pad for immediate recovery. ABRs were measured using a custom acoustic system (Eaton-Peabody Laboratories, EPL, Massachusetts Eye and Ear Infirmary, MEEI) in an electrically shielded and sound attenuating chamber. All recordings were performed with the researcher blinded to genotype.

ABRs were recorded from three subcutaneous needle electrodes: a recording electrode caudal to the pinna, a reference electrode at the vertex, and a ground electrode by the tail. ABR stimuli were typically presented from 20 to 80 dB SPL in 5 dB steps at frequencies specified in each ABR graph. Each stimulus was presented as a 5 ms tone-pip at a rate of 31/s, with a 0.5 ms rise-fall time and alternating polarities. Responses were amplified 10,000x, filtered with a 0.3–3 kHz passband (P511, Grass), and averaged 512 times. Recordings with peak-to-peak amplitudes exceeding 15 μV were rejected as artifacts. ABR thresholds and amplitudes were analyzed using ABR Peak Analysis software (v1.9.0; EPL MEEI Engineering Core). ABR threshold was defined as the lowest stimulus level at which wave-I could be identified at a consistent latency by qualitative visual inspection. If animals showed no response at 80 dB SPL, a threshold of 85 dB was assigned.

### Immunohistochemistry

No more than three days after the final ABR, animals were transcardially perfused and cochlea post-fixed with 4% paraformaldehyde (PFA) for 2 hours at room temperature, followed by overnight fixation at 4°C. Tissues were then rinsed three times with 1X phosphate-buffered saline (PBS) for 5 minutes each to remove residual PFA. Decalcification was performed in 120 mM ethylenediaminetetraacetic acid (EDTA) at room temperature on a shaker for three days. After decalcification, tissues were rinsed again with 1X PBS (3 × 5 min), cryoprotected overnight in 30% sucrose at 4°C, and transferred to −80°C for storage. Cochleae were thawed in room temperature water and rinsed in 1X PBS (3 × 5 min) to remove sucrose, and micro-dissected in 1X PBS. Cochlear turns were transferred to a blocking solution (5% normal donkey serum and 1% Triton X-100) for 1 hour at room temperature on shaker, followed by incubation in a Fab blocking solution (1:10 Donkey anti-Mouse IgG Fab fragment, 1% Triton X-100) for 30 minutes. Primary antibodies were applied in 1% serum and 0.3% Triton X-100 and incubated overnight at 37°C on a shaker. The following antibodies were used: anti-Calb2 (1:500, Swant CG1), anti-CTBP2 (1:500, BD Transduction Laboratories), anti-Homer1 (1:500, SynapticSystems), and anti-dsRed (1:500, Takara). Samples were incubated with secondary antibodies in 1% serum and 0.3% Triton X-100 for 2 hours at 37°C, with the solution replaced by a fresh identical secondary antibody solution at 1 hour. Rinses after primary and secondary antibody were done using 1% PBST for 10 minutes at room temperature.

### Image acquisition

All images were acquired on a Zeiss LSM 800 confocal microscope. High-magnification z-stack images were acquired using 63× oil-immersion objective. Laser power and digital gain were adjusted for each stack to maximize signal-to-noise-ratio while minimizing the number of saturated pixels with researcher blind to genotype and exposure condition. Cochlear frequency maps were generated by tracing the entire length of the cochlear spiral in 10X images of whole mounts using a custom ImageJ plugin available through the Histology Core at MEEI (https://meeeplfiles.partners.org/Measure_line.class) that translates cochlear position into frequency according to the published map for the mouse^56,57^. On rare occasions, cochlear turns that did not contain the 32 kHz region were lost during immunolabelling. In these cases, a surrogate micrograph that contained a cochlear turn of a similar length and always from the same region were used to aid in mapping. If the missing turn contained the 32 kHz region, as is easy to approximate based on morphology of the tissue, then the sample was omitted from analysis.

### Synapse and Terminal Analysis

Synaptic and terminal status for all markers was performed as described in Supplementary Fig. 2 and in the accompanying GitHub repository. Following high magnification imaging, pre- and post-synaptic markers were reconstructed in Imaris and all surface statistics were exported using default software features. Automatic thresholding was used for the generation of all surfaces, though the final threshold value used was manually adjusted for each image to ensure that all markers within the CALB2 IHC boundary were captured. For *Ntng1*^*Cre*^*;Ai14* volume analysis, these surfaces were manually refined to remove artefactual volume that often occurred as a result of puncta being merged during surface creation or surface enlargement that occurred due to spherical aberrations, which are more common when imaging in the more basal regions of the sensory epithelium. Surface statistics were then converted into Python Pandas format using custom code (Jupyter Lab notebooks), and then imported to the custom SynAnalyzer ImageJ code for thumbnail array generation and manual scoring. In training sessions prior to scoring, users were instructed to use the distribution of any tdTomato signal across the field of view within a given thumbnail to score overlap with the Homer1 punctum in question independently of signal intensity. All downstream analysis was subsequently performed in Python notebooks using custom code to compile measures from all images and perform additional image-specific quantification of parameters (e.g., synaptic index, orphan index, tdTomato+ synapse proportion, normalized volume). All code will be made freely available for future use via the Goodrich Lab GitHub repository at the time of publication.

### Quantification of puncta properties

For all reported indices (i.e., Synaptic Index, Ribbon Index, and Orphan Index), the respective value was calculated for each field of view (one per animal) by taking the total number of puncta satisfying the criteria in question (e.g., synaptic Homer1 puncta) for that field of view, and dividing it by the number of IHCs analyzed for surface reconstructions in that field of view. All calculations were performed in custom Python-based Jupyter Lab notebooks as follows. Homer1 volumes were filtered to include only those belonging to control animals for a given experiment and for those belonging to a synaptic pair. Then, the median volume within this subset was identified. Finally, all Homer1 volumes (both synaptic and orphan) were divided by this control median volume.

## Statistics

GraphPad Software was used to perform ABR-related statistical analyses (Prism GraphPad version 10.5.0). For analysis of wave-I amplitudes, nonlinear regression (NLR) analysis was performed to determine whether both groups are fit by the same 2nd-order polynomial equation or required separate equations. In this analysis, initial value (“B0”) was constrained to equal zero. Error bars were limited to a minimum range of zero uV. The null hypothesis (i.e., “one curve for all data sets”) was rejected if alpha<0.05, and the alternative hypothesis (i.e., “different curve for at least one data set”) was accepted. The Python Scipy.Stats package was used for all other analyses. A description of the exact test used for each comparison is provided in the corresponding figure legend.

## Acknowledgments

The authors thank Goodrich Lab members for their assistance in manuscript review and editing. They thank Lucy Lee for guidance on synapse reconstruction methods, Dr. Gabriel Romero on ABR and Wave-I statistical analysis, and Dr. Charlie Liberman for his guidance on synapse immunolabelling methods and noise exposure parameters, and the Xu-Friedman lab (University of Buffalo) for shipping *Calb2*^*CreERT2*^ and *Lypd1*^*CreERT2*^ mice. The research and publication of this work was supported by a Blavatnik Sensory Disorders Research Grant awarded to LVG, and by the National Institute on Deafness and Other Communication Disorders of the National Institutes of Health under Award Number R01DC009223 to LVG. JAF was supported by funding from the National Institute on Aging of the National Institutes of Health under Post-doctoral Transition Award Number K00AG078230.

## Author contributions

This project was conceptualized by LVG, TGC, and JAF, who collectively established the methodology, designed the experiments, and analyzed results. JAF developed custom Python and ImageJ code for image analysis. RDM assisted JAF and TGC with data curation and experimental execution. LVG and JAF oversaw project administration and wrote the original draft of the manuscript which was reviewed and edited by all authors. LVG supervised the study and acquired funding.

