## Supplemental Figures 1 to 5 and Table 1 for "Molecularly defined auditory neuron subtypes show different vulnerabilities to noise- and age-related synaptopathy in mice"

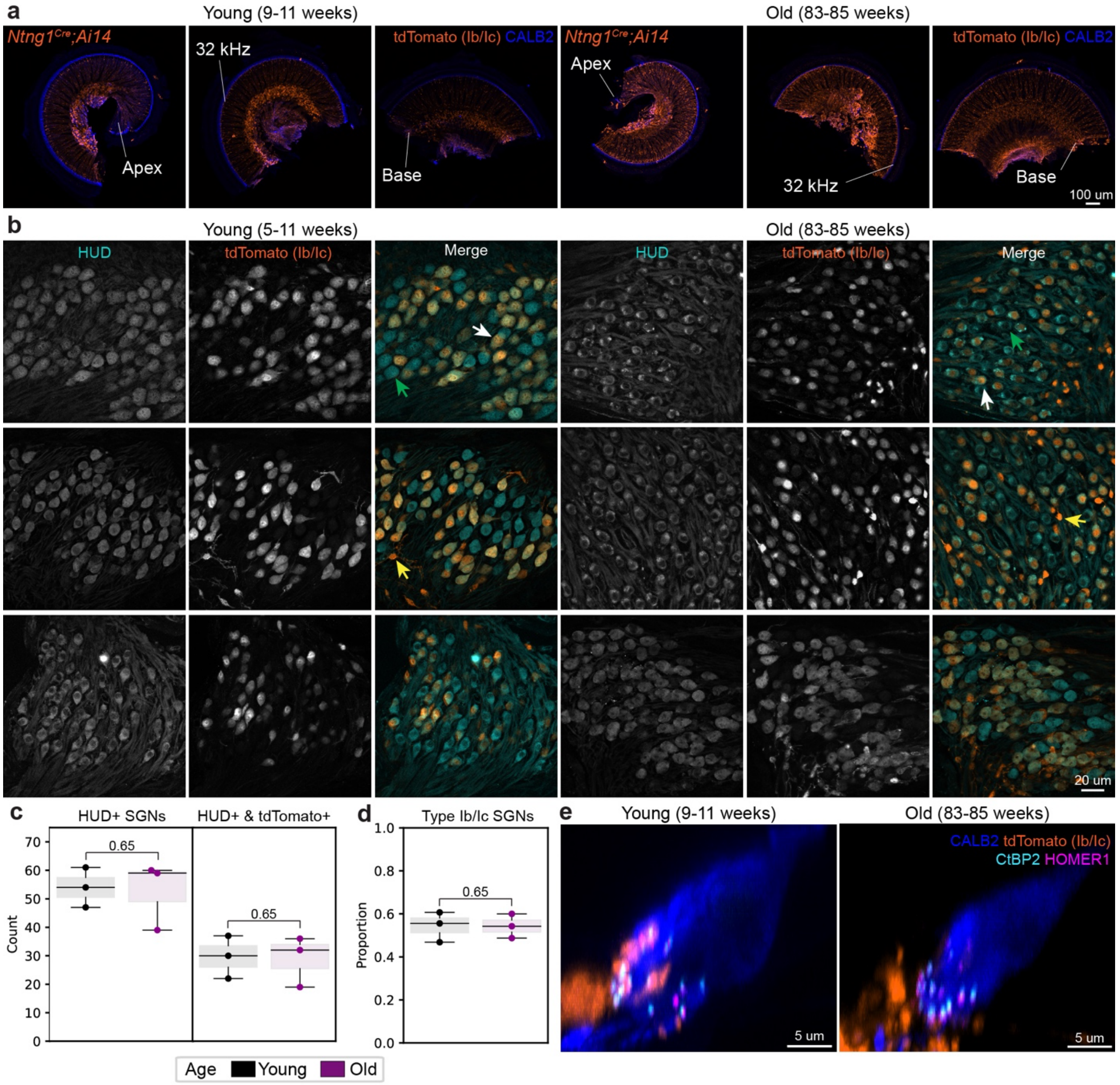


**Supplemental Fig. 1: Type Ib/Ic SGNs are labelled with tdTomato regardless of age. a** Low-power micrographs of cochlear turns from *Ntng1*$^{Cre}$*;Ai14* animals used for aging analysis show tdTomato expression SGN somata at all tonotopic regions at both young and old ages. **b** High-power confocal images of cochlear sections immunolabelled for HUD and tdTomato used to quantify the proportion of Ib/Ic SGNs. **c** SGN counts for all images analyzed showing the number of HUD+ somata for each image (one field of view per animal), and the number of HUD+/tdTomato+ SGNs for each image. **d** Relative proportion of Type Ib/Ic SGNs in each field of view based on quantification shown in **c**. For **c** and **d**, each data point represents a single field of view (one field of view per animal, young N=3, old=3), boxes represent interquartile ranges (IQRs), horizontal lines represent the median, box height represents the IQR, and whiskers extend to data points farthest from the median but within 1.5x the IQR from the box edge; one-sided Mann-Whitney test used for all statistics. **e** Maximum-intensity projection images of virtual slices showing the inclusion of the CALB2 channel, corresponding to Fig. 1. (Note: the IHC shown for an old animal is tilted into the page and thus appears much smaller in volume. IHC size qualitatively appears more variable in older animals, but we did not observe a consistent bias towards larger or smaller IHCs relative to young controls.)

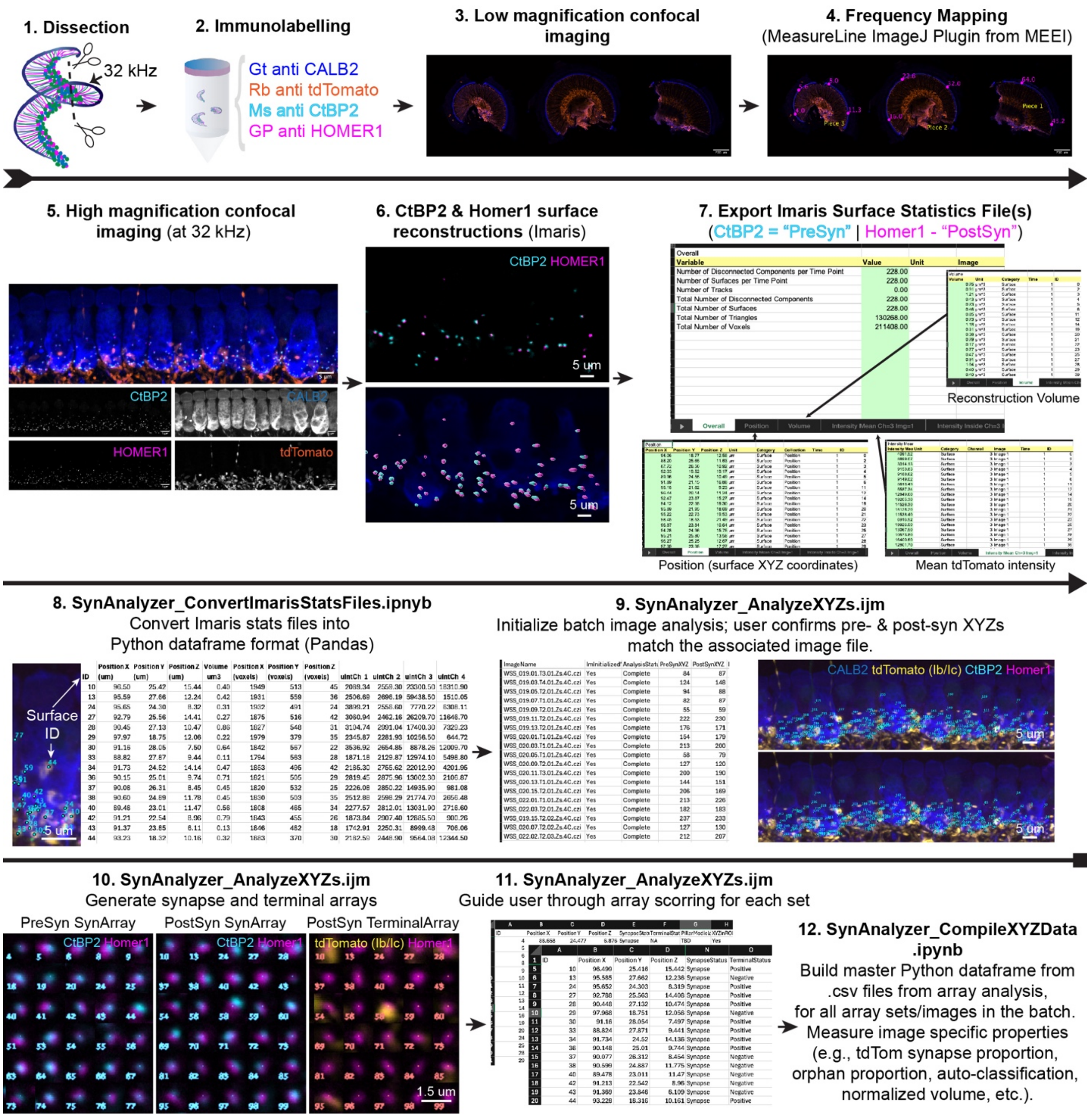


**Supplemental Fig. 2: Pipeline for subtype-specific analysis of synaptic markers in high-power micrographs. Steps 1-5**: Tissue dissection, immunolabelling, low-power imaging, tonotopic frequency mapping to identify the 32 kHz region of the dissected tissue, and high-power imaging of the tissue at the 32 kHz region. **Steps 6-7**: Imaris based reconstruction of pre- and post-synaptic markers, and export of all surface statistics using default Imaris tools. **Steps 8-11:** Custom Python and ImageJ based analysis of high-power micrographs based on data exported from Imaris. An unique surface ID is assigned to every reconstructed punctum. IDs are tracked throughout the pipeline along with their "pre-" or "post-" synaptic identity and the image ID to prevent the overloading of IDs. Users first confirm that the underlying dataset for the reconstructions corresponds to the appropriate micrograph and are in the appropriate position prior to thumbnail generation. Thumbnails for each of the synaptic marker channels and the tdTomato channel are generated for every reconstructed surface and assembled into synaptic or terminal arrays for user scoring based on the overlap of tdTomato signal with the Homer1 signal, regardless of signal intensity. The custom ImageJ macro walks the user through the process of scoring each array set. In training sessions prior to analysis, users are instructed to consider the distribution of the tdTomato signal across the entire thumbnail in order to differentiate uniform background signal from instances were tdTomato levels are low but spatially restricted to an apparent terminal that overlaps with the Homer1 punctum in question. **Step 12**: Files from the entire batch image analysis (one per synaptic marker array set per animal) are compiled using custom Python code where subsequent calculations of metrics such as synaptic index, tdTomato+ synapse proportion, and normalized volumes are calculated for the thousands of synaptic markers belonging to each batch, where one batch corresponds to a single experiment (e.g., Aging).

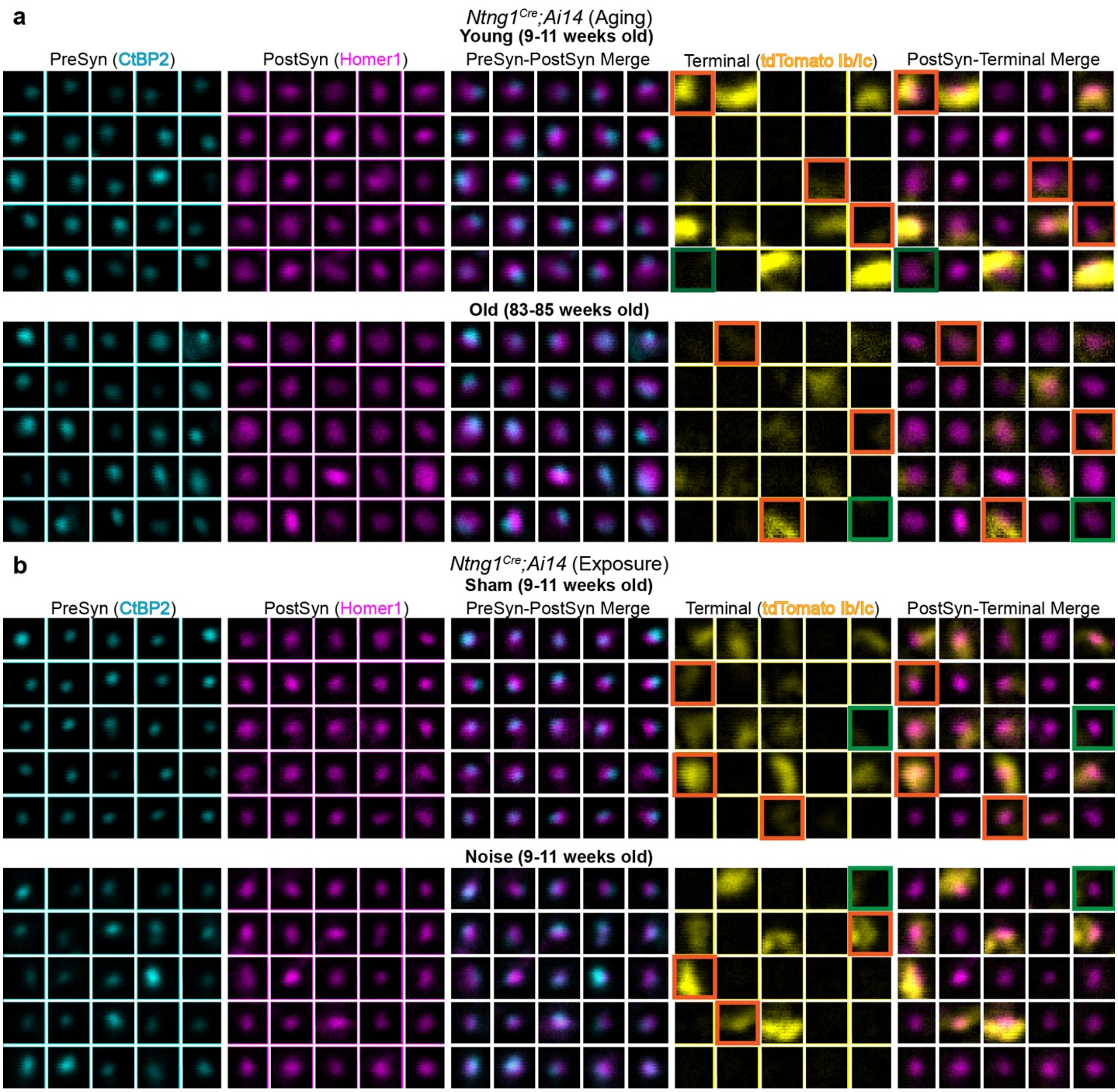


**Supplemental Fig. 3: Representative synaptic and terminal thumbnails generated by SynAnalyzer.** Thumbnail arrays generated by custom ImageJ macro, SynAnalyzer, of pre- and post-synaptic markers and tdTomato terminals for all conditions studied in *Ntng1$^{Cre}$;Ai14* mice. Thumbnails were randomly selected from a single high-power image for each condition after orphaned puncta were excluded from the array set. Orange squares highlight tdTomato+ terminals. Highlighted examples show the range of intensities analyzed for each condition. Green squares show thumbnails that have tdTomato signal within the square but are tdTomato- since the signal does not overlap with the associated Homer1 punctum.

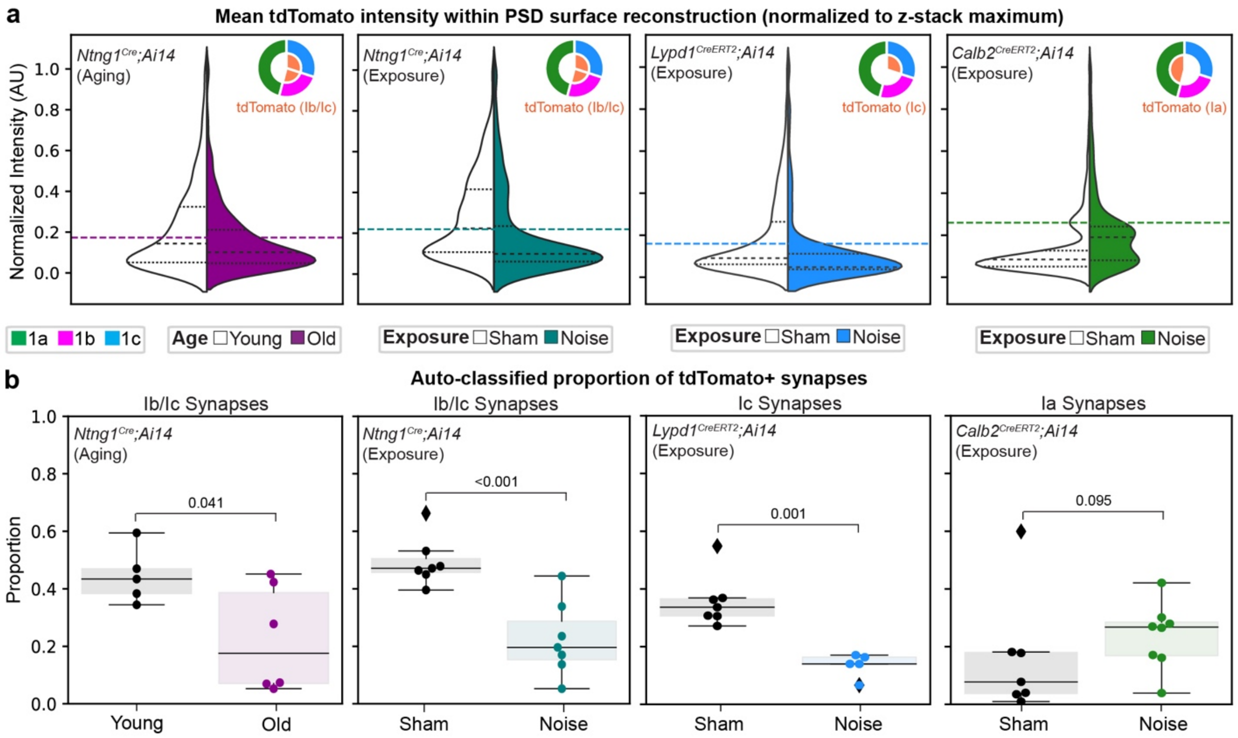


**Supplemental Fig. 4: Automated analysis of molecular subtype synapse proportions. a** Split violin plots of mean tdTomato intensity within each Homer1 surface reconstruction, as calculated by Imaris and normalized to the z-stack maximum, for only those Homer1 puncta that belong to a synapse. Horizontal lines indicate the tdTomato intensity used as a cutoff for classifying tdTomato status for each experiment data set. These values are calculated in Python based on the population statistics from the control dataset for that experiment. For example, in the case of *Ntng1$^{Cre}$;Ai14* aging experiments, 55% of synapses in young animals were scored as being tdTomato-. Using the Python Numpy "percentile" function, the 55th percentile of tdTomato intensities within each of the synaptic Homer1 surface reconstructions was calculated as 0.18. This value was then used as the cutoff for that experiment's dataset where Homer1 surfaces with a tdTomato intensity below 0.18 were classified as tdTomato- and those above as tdTomato+. **b** Quantification of the molecular subtype synapse proportion for each condition and experiment based on the automated classification of tdTomato status described in **a**. An exclusion criterion was established at the beginning of the analysis such that any animals having more than 90% of synapses being classified as tdTomato+ would not be included. This resulted in the exclusion of one animal for the aging dataset and one animal from the *Calb2$^{CreERT2}$;Ai14*. Each data point represents a single field of view (one field of view per animal) boxes represent interquartile range (IQRs), horizontal lines represent the median, box height represents the IQR, and whiskers extend to data points farthest from the median but within 1.5x the IQR from the box edge, diamonds mark data points outside whiskers and are classified as potential outliers by Python Seaborn package; one-sided Mann-Whitney test used for all statistics and include data points classified as potential outliers.

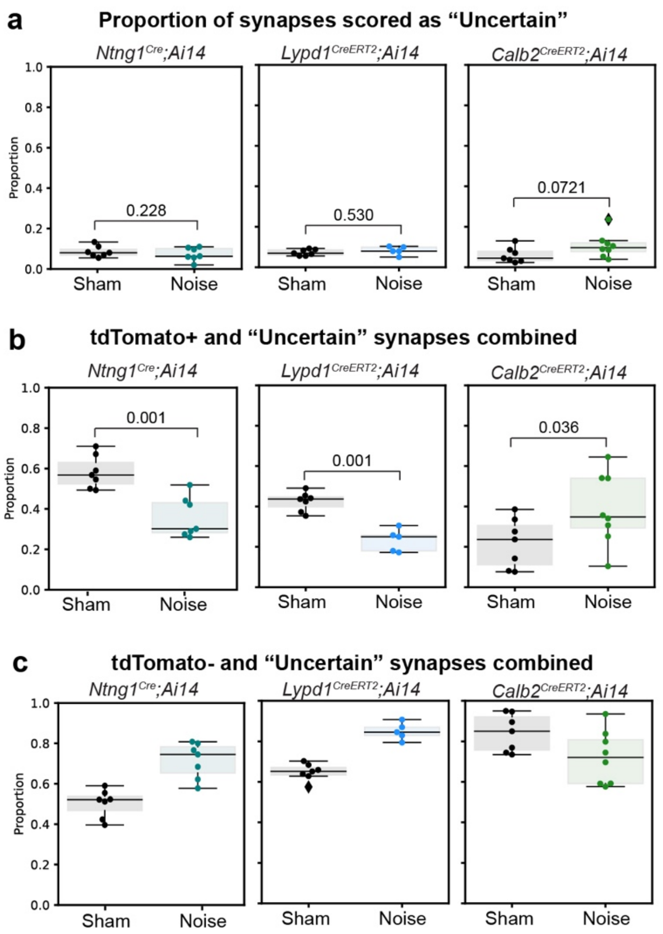


**Supplemental Fig. 5: Analysis of synapses scored as having “uncertain” tdTomato status. a** Proportion of synapses scored as “uncertain” for all experiments. Proportions do not significantly vary with noise exposure for any group. **b** Treating all synapses scored as “uncertain” as if they are tdTomato+ does not alter the outcomes of tdTomato+ alone analysis. **c** Combining all “uncertain” synapses with tdTomato- synapses is complementary to the results in **b**, demonstrating that these three synapse statuses represent the complete pool of synapses. For all panels, individual data points correspond to the same fields of view shown in main figures for each experiment. Each data point represents a single field of view (one field of view per animal) boxes represent interquartile range (IQRs), horizontal lines represent the median, box height represents the IQR, and whiskers extend to data points farthest from the median but within 1.5x the IQR from the box edge, diamonds mark data points outside whiskers and are classified as potential outliers by Python Seaborn package; one-sided Mann-Whitney test used for all statistics and include data points classified as potential outliers.

**Supplementary Table 1.** ABR Thresholds for *Lypd1$^{CreERT2}$;Ai14* and *Calb2$^{CreERT2}$;Ai14* mice

| **Condition** | **Timepoint** (post-exposure) | **Reporter Line** | **N** | **8 kHz** (mean+/-SEM) | **16 kHz** (mean+/-SEM) | **32 kHz** (mean+/-SEM) |
|---|---|---|---|---|---|---|
| Sham | One day | *Lypd1$^{CreERT2}$;Ai14* | 7 | 32.14+/-1.49 | 20.71+/-0.71 | 27.86+/-2.40 |
| | | *Calb2$^{CreERT2}$;Ai14* | 8 | 34.38+/-1.75 | 21.25+/-0.82 | 36.25+/-2.27 |
| | One week | *Lypd1$^{CreERT2}$;Ai14* | 7 | 30.71+/-1.30 | 21.43+/-0.92 | 30.71+/-1.30 |
| | | *Calb2$^{CreERT2}$;Ai14* | 8 | 32.50+-/0.94 | 21.88+/-1.32 | 36.25+/-1.83 |
| Noise | One day | *Lypd1$^{CreERT2}$;Ai14* | 6 | 37.50+/-2.81 | 44.17+/-4.90 | 73.33+/3.07 |
| | | *Calb2$^{CreERT2}$;Ai14* | 8 | 35.63+/-1.75 | 50.71+/-3.52 | 73.13+/-2.82 |
| | One week | *Lypd1$^{CreERT2}$;Ai14* | 6 | 32.50+/-2.14 | 23.33+/-2.11 | 39.17+/-2.01 |
| | | *Calb2$^{CreERT2}$;Ai14* | 8 | 31.88+/-1.32 | 27.50+/-3.27 | 35.00+/-1.64 |
